# Lac1-dependent ceramide synthesis couples septum assembly to cell separation in *Schizosaccharomyces pombe*

**DOI:** 10.64898/2026.10.31.763146

**Authors:** Ana Serrano, Laura Gómez-Delgado, Kazuki Hanaoka, Kuya Matsunaga, Kouichi Funato, Juan Carlos G. Cortés, Juan Carlos Ribas, María Alcaide-Gavilan, Ignacio Flor-Parra, Rafael Lucena

## Abstract

Ceramides are precursors of complex sphingolipids and bioactive membrane lipids. In the fission yeast *Schizosaccharomyces pombe*, they are produced by a ceramide synthase with two catalytic subunits, Lac1 and Lag1, which are redundant but not functionally equivalent: only Lac1 is required for normal growth and cell size. Why Lac1 has this unique role, and how ceramide synthesis contributes to cell morphogenesis, remain unknown. Here, we show that Lac1 couples septum assembly to cell separation. Cells lacking Lac1 die predominantly during cell separation and build septa with a thinner primary layer and markedly thicker secondary layers, accompanied by altered deposition of β-(1,3)-glucans and galactomannan. These defects are not rescued by osmotic stabilization. The essential glucan synthase Bgs4 is mislocalized to cytoplasmic granules, and total β-(1,3)-glucan synthase activity is partially reduced. Nevertheless, Ags1, Bgs1 and Bgs4 accumulate to higher levels and persist almost twice as long at the division site, indicating defective clearance rather than delivery of the septum-synthesis machinery. Plasma membrane sterol distribution remains unaffected. To accurately impair ceramide synthesis, we isolated thermosensitive *lac1-ts* alleles in a *lag1*Δ background. Their substitutions cluster in the conserved TRAM/LAG1 domain. Metabolic labeling showed that these alleles reduce ceramide synthesis at the restrictive temperature. Molecular dynamics simulations indicated that two substitutions perturb distinct contacts with C26-CoA, while Lac1-M167T loses its endoplasmic reticulum localization. Acute inactivation recapitulates the deletion phenotype, causing septum accumulation and cell death. Together, our findings establish that Lac1-dependent ceramide synthesis is required for the orderly completion of septum assembly and cell separation.

## INTRODUCTION

Regulation of cell growth is essential for the survival and virulence of fungal pathogens [1]. Unlike animal cells, fungi are surrounded by a rigid cell wall of polysaccharides that provides mechanical support and protects the cell from environmental stresses [2]. This structural constraint creates a fundamental challenge: cell growth and division require the coordinated expansion of the plasma membrane with the synthesis, remodeling, and deposition of new cell wall material. Failure to maintain this coordination can result in cell lysis or defects in cytokinesis [3]. Thus, fungal cells must precisely coordinate plasma membrane growth with cell wall biogenesis to maintain the cellular integrity throughout the cell cycle [4,5]. Although several signaling pathways that regulate fungal growth have been identified, how information about membrane growth is coupled to the spatial regulation of cell wall synthesis remains poorly understood [5–8].

The fission yeast *Schizosaccharomyces pombe* provides an established model to investigate the mechanisms that coordinate cell growth and division. *S. pombe* grows primarily through polarized extension of its cell ends during interphase and divides through the formation and subsequent cleavage of a medial division septum during cytokinesis [9]. Septum formation is a highly coordinated process requiring the synthesis and remodeling of cell wall polymers together with plasma membrane expansion and actomyosin ring constriction [10]. Septation is also highly coordinated with the nuclear mitosis, starting the formation of a septum rudiment during early anaphase B, when the two nuclei start to separate [11]. Unlike animal cells, in which the cleavage furrow ingresses by constriction of the actomyosin ring alone, fungi build a septum wall in the wake of the ring, and the two processes are mechanically interdependent: septum synthesis sets the rate at which the ring constricts, while the ring spatially guides septum growth and keeps its leading-edge circular [12–15]. Septation is therefore an integral part of cytokinesis in *S. pombe* rather than a subsequent event. Cell division is completed only after the closed septum has matured and its internal layer of primary septum, together with the surrounding septum edging, is digested by the Eng1 and Agn1 glucanases, allowing the two daughter cells to separate [16].

The *S. pombe* cell wall consists of three ultrastructural layers, with electron-dense outer and inner layers containing galactomannan surrounding a central layer enriched in α-glucans, β-(1,3)-glucans, and β-(1,6)-glucans [17]. These polymers are synthesized by a set of glucan synthases, including the essential enzymes Ags1 [18,19], Bgs1 [20,21], Bgs3 [22], and Bgs4 [16,17], whereas galactomannan synthesis involves multiple non-essential proteins [23]. The organization and relative contribution of these polymers provide the cell wall with the mechanical properties required to resist turgor pressure while allowing controlled remodeling during growth and division [13]. Despite detailed knowledge of cell wall composition, how the synthesis and spatial organization of these components are coordinated with plasma membrane growth remains poorly understood [3,14,24,25]. Because the fungal cell wall is both essential for fungal viability and absent from mammalian cells, pathways controlling its synthesis and organization represent attractive targets for antifungal intervention [26,27].

Membrane lipids are increasingly recognized not only as structural components but also as important regulators of membrane organization and signaling [28–30]. Among these, sphingolipids have emerged as critical regulators of fungal growth, membrane homeostasis, and cell integrity [31–34]. Ceramides occupy a central position in sphingolipid metabolism, serving both as precursors for complex sphingolipids and as bioactive membrane lipids capable of influencing membrane properties and protein organization [35–37]. Their long, saturated acyl chains promote tight lipid packing and can influence the organization and physical properties of membrane domains [35]. Alterations in ceramide metabolism have been associated with diverse cellular processes, including proliferation, stress responses, and host–pathogen interactions [36,38–42]. However, whether ceramide synthesis contributes directly to the spatial organization of fungal cell wall synthesis remains largely unexplored.

Sphingolipid biosynthesis is highly conserved from yeast to humans. In the endoplasmic reticulum (ER), serine palmitoyltransferase catalyzes the condensation of serine and palmitoyl-CoA to generate 3-ketodihydrosphingosine, which is subsequently reduced to the long-chain base dihydrosphingosine (DHS) [43]. DHS can be hydroxylated by Sur2 to generate phytosphingosine (PHS) [44]. These long-chain bases are then N-acylated with very-long-chain fatty acids (VLCFAs) by the ceramide synthase (CerS) complex to produce ceramides. In yeast, CerS enzymes preferentially use VLCFAs, predominantly C_24_ and C_26_, generating ceramides with exceptionally long acyl chains that contribute to membrane organization and function [45].

In *S. pombe*, ceramide synthesis is mediated by the two catalytic CerS subunits Lac1 and Lag1, which are functionally redundant but essential, and were recently characterized by our group [46]. Newly synthesized ceramides are transported from the ER to the Golgi, where they serve as precursors for complex sphingolipids, including inositol-phosphoceramide (IPC) and mannosyl-IPC (MIPC). Unlike *S. cerevisiae*, *S. pombe* does not synthesize M(IP)₂C, making MIPC the terminal and most abundant complex sphingolipid in this organism [47]. Thus, Lac1 and Lag1 occupy a central position in the pathway linking VLCFA metabolism to the production of fungal sphingolipids.

Our previous work in fission and budding yeast demonstrated that ceramide synthesis has a previously unrecognized role in regulating cell growth and size homeostasis [34,46,48]. Importantly, in fission yeast this function is not equally shared by the two CerS catalytic subunits. Loss of Lac1, but not Lag1, causes a pronounced reduction in growth rate and defects in cell-size control [46]. These findings revealed a functional divergence between Lac1 and Lag1 and identified Lac1 as a key regulator of growth-associated ceramide metabolism in *S. pombe*. However, the cellular processes responsible for the distinct requirements for Lac1 remained unclear.

Here, we investigated whether the requirements for Lac1 reflect a role for ceramide synthesis in maintaining cell envelope integrity and coordinating cell wall assembly with cytokinesis. We have found that cells lacking Lac1 die during cell separation and assemble division septa with an abnormally thick layered architecture. Moreover, the glucan synthases that build the septum are mislocalized and abnormally retained at the division site, while the distribution of plasma membrane sterols is unaffected. Because deletion of both ceramide synthases is lethal, and because the *lac1*Δ phenotype may be shaped by chronic adaptation and compensation with the Lag1 function, we isolated thermosensitive Lac1 alleles in a *lag1*Δ background. These alleles lower ceramide synthesis at the restrictive temperature and reproduce the septation defect and acute cell death. Together, our results connect Lac1-dependent ceramide synthesis to the spatial and temporal control of cell wall and septum synthesis, and provide a framework for understanding how membrane lipid metabolism contributes to the maintenance of cell integrity during fungal cell growth and division.

## RESULTS

### Ceramide synthase Lac1 is essential for maintaining cell wall integrity and septum organization

In *Schizosaccharomyces pombe*, ceramides are synthesized at the endoplasmic reticulum (ER) by the ceramide synthase subunits Lac1 and Lag1 [49] (Dawson et al., 2008). Our previous work established that loss of Lac1, but not Lag1, causes pronounced defects in cell growth and alters sphingolipid composition, suggesting that these two ceramide synthase subunits have distinct cellular functions [46].

To investigate the consequences of Lac1 loss for cell viability, we performed fluorescence microscopy of *lac1*Δ and *lag1*Δ cells stained with the fluorescent dye calcofluor white, which preferentially stains linear β-(1,3)-glucan. Strikingly, cell death occurred predominantly during cytokinesis immediately following septum degradation and cell separation. Specifically, 26.26% of *lac1*Δ cells (n=1,112) underwent catastrophic cell death during cell separation, compared with 1.70% of wild-type cells (n=1,179) and 3.33% of *lag1*Δ cells (n=1,110) (**Figure 1A**). The strong association between Lac1 loss and cell death during cell separation suggested that the primary defect may involve septum formation, septum organization, or cell wall remodeling during cytokinesis.

**Figure 1.**
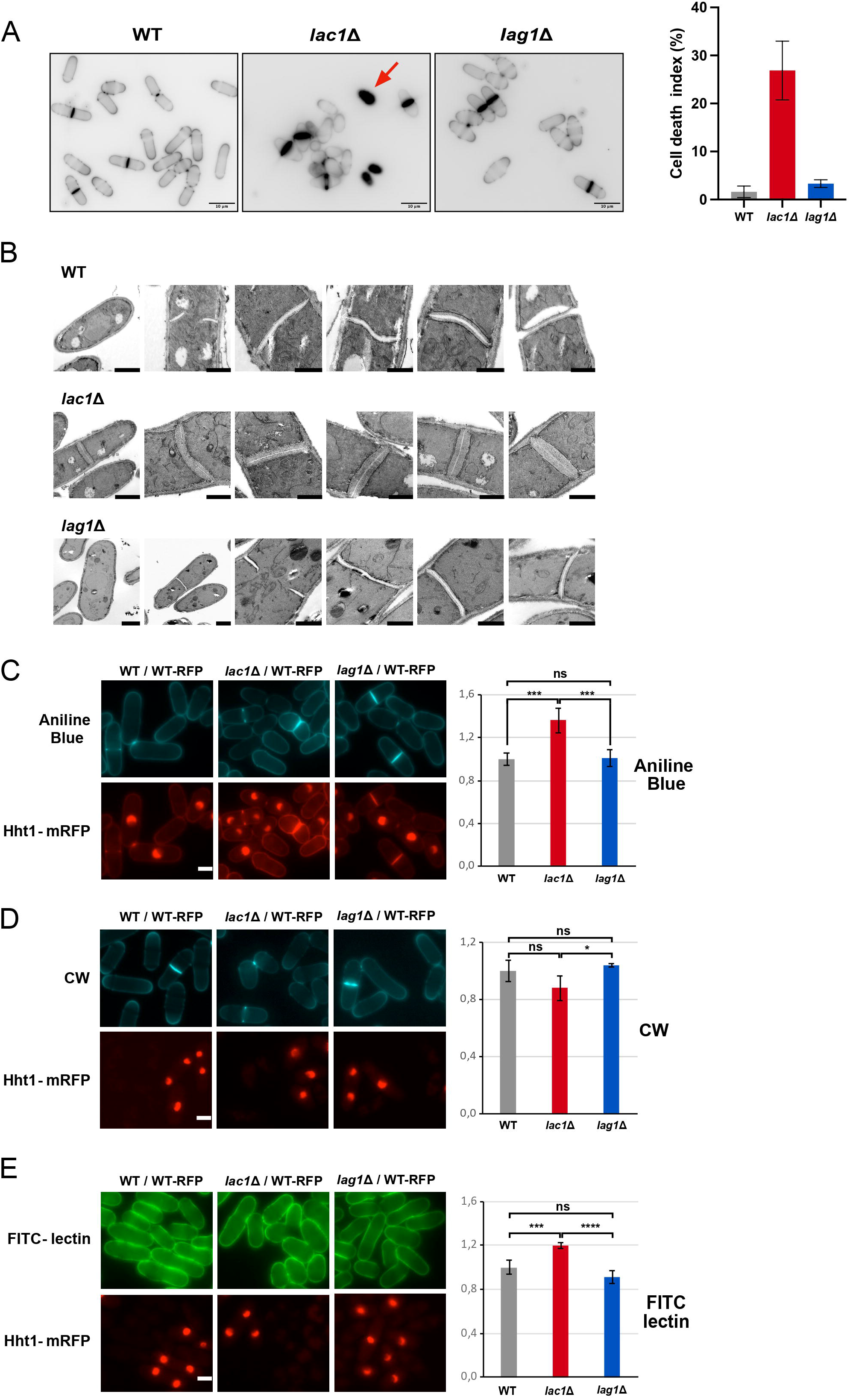
Loss of Lac1 causes cell death during cell separation and alters septum and cell wall polymers deposition. **(A)** Cell death in wild-type, *lac1Δ* and *lag1Δ* cells. Left, representative calcofluor white images; cells scored as dead showed intense calcofluor staining (red arrow). Scale bar, 10 µm. Right, cell death index, scored as the percentage of cells undergoing death during or immediately after cell separation (n = 1,179 wild-type, 1,112 *lac1Δ* and 1,110 *lag1Δ* cells, from 3 independent experiments). Bars show the mean ± SD **(B)** Transmission electron micrographs of wild-type, *lac1Δ* and *lag1Δ* cells. Left, entire cells; right, division septa at higher magnification. Wild-type and *lag1Δ* cells display the characteristic three-layered septum, with an electron-transparent primary septum flanked by electron-dense secondary septa, whereas *lac1Δ* cells show a thinner primary septum and markedly thicker secondary septa. Scale bars, 2 µm (entire cells) and 1 µm (septum details). **(C–E)** Cell wall polymers staining. Each strain was mixed with an Hht1-mRFP-expressing wild-type (WT) reference strain and imaged in the same field (WT/WT-RFP, *lac1Δ/*WT-RFP, *lag1Δ*/WT-RFP); histone Hht1-mRFP identifies the reference cells, and fluorescence intensities were normalized to the internal control within WT-RFP within each field. **(C)** Aniline blue, which detects β-(1,3)-glucans at the cell wall, and primary and secondary septum; **(D)** Calcofluor white (CW), which detects linear β-(1,3)-glucan mainly at the primary septum and in lower amount at the cell wall; **(E)** FITC-conjugated lectin from *Bandeiraea simplicifolia*, which detects the galactomannan of the cell wall surface. Left, representative images of each staining together with the Hht1-mRFP channel. Scale bars, 5 µm. Right, quantification of mean fluorescence intensity in the cell wall normalized to the wild-type (WT-RFP) value, set to 1 (number of cells from 7 independent experiments: Anilin blue: *lac1*Δ/WT: 121/139; *lag1*Δ/WT: 93/95; WT/WT: 78/94. CW: *lac1*Δ/WT: 105/98; *lag1*Δ/WT: 89/89; WT/WT: 153/89. FITC-lectin: *lac1*Δ/WT: 125/73; *lag1*Δ/WT: 118/89; WT/WT: 106/101). Bars show the mean ± SD; statistical significance was assessed by one-way ANOVA, followed by Tukey’s post hoc test (ns, not significant; *, p < 0.05; ***, p < 0.001; ****, p < 0.0001).

Because defects in cell wall integrity can increase susceptibility to turgor pressure and osmotic lysis, we next tested whether osmotic stabilization could alleviate the cell growth defect of *lac1*Δ cells. Serial dilution spot assays performed in the presence of 1.2 M sorbitol failed to rescue even partially the growth defect of *lac1*Δ cells, and showed to be slightly harmful (**Supplementary Figure 1**). Thus, the phenotype associated with Lac1 loss cannot be attributed to a simple increase in osmotic fragility that can be bypassed by external osmotic support, suggesting a broader defect in cellular integrity or morphogenesis.

We next examined whether Lac1 loss affects the ultrastructure of the cell wall and division septum using transmission electron microscopy (TEM). Wild-type and *lag1*Δ cells displayed the characteristic three-layered division septum, consisting of an electron-transparent primary septum flanked by electron-dense secondary septa on both sides. Both strains exhibited similar septum thickness, and also relatively uniform cell wall architecture, although slightly thinner in *lag1*Δ cells. In contrast, *lac1*Δ cells showed a pronounced alteration in septum ultrastructure. Their division septa were overall substantially thicker than those of wild-type and *lag1*Δ cells, with a thinner electron-transparent primary septum and markedly thicker electron-dense secondary septa. Despite these alterations in septum organization, the general cell wall architecture of *lac1*Δ cells remained relatively uniform (**Figure 1B**).

To determine whether these ultrastructural abnormalities were accompanied by alterations in the deposition of specific cell wall polysaccharides, we analyzed major cell wall components using fluorochromes (aniline blue and calcofluor white), and a fluorophore-conjugated probe (FITC-lectin). To control for variability in staining and illumination, each strain was mixed with a nuclear histone Hht1-mRFP-marked wild-type reference strain and imaged in the same field, and fluorescence intensities measured in the cell poles were normalized to these of the internal wild-type control. Aniline blue staining, which detects β-(1,3)-glucans associated with the cell wall and both the primary and secondary septa, revealed increased fluorescence intensity in *lac1*Δ cells (136%) compared with both wild-type (100%) and *lag1*Δ cells (101%), whereas wild-type and *lag1*Δ cells did not differ (**Figure 1C**). Next, we analyzed the fluorescence of calcofluor white staining. This fluorochrome binds specifically and with high affinity to the linear β-(1,3)-glucan located mainly in the primary septum, and in a small amount in the cell wall [21]. *lac1Δ* cells showed a consistent decrease in fluorescence intensity (88 %) that did not reach significance relative to wild-type (100%) but was significant relative to *lag1Δ* cells (104 %); this reduction of fluorescence in *lac1Δ* cell walls was consistent with the thinner primary septum of *lac1Δ* cells observed by TEM (**Figure 1B, D**). Finally, we examined the galactomannan-containing outer matrix layer of the cell wall using FITC-conjugated lectin from *Bandeiraea simplicifolia* (*Griffonia simplicifolia*), which binds specifically to the galactose residues of the galactomannoproteins located in the cell wall surface. *lac1Δ* cells exhibited increased fluorescence (120 %) compared with both wild-type (100%) and *lag1Δ* cells (91 %), suggesting altered accumulation or exposure of galactomannan-associated epitopes at the cell wall surface, while wild-type and *lag1Δ* cells were again indistinguishable (**Figure 1E**). For all three probes (fluorochromes and FITC-lectin), therefore, cell wall polymer deposition was altered specifically by the loss of Lac1, with *lag1Δ* cells remaining indistinguishable from wild-type (**Figure 1C-E**). To analyze whether the alterations observed in *lac1Δ* were also present in the septum walls, aniline blue and calcofluor fluorescence of septa of wild-type, *lag1Δ*, and *lac1Δ* cells were examined by scanning a line along the entire septum length **(Supplementary Figure 2**). Like in the cell walls, *lac1*Δ cells showed a similar increase in aniline blue fluorescence intensity compared with that of wild-type, being more apparent in the middle region of the septum, whereas no differences were detected with calcofluor, suggesting that although *lac1*Δ cells contain a thinner primary septum, the concentration of total β-(1,3)-glucan in the primary septum is increased whereas that of linear β-(1,3)-glucan remains similar to that of wild-type. Similarly, no differences were detected in the septum fluorescence with either aniline blue o calcofluor of *lag1*Δ cells compared to these of wild-type, or between wild-type control cells with and without Hht1-mRFP **(Supplementary Figure 2**).

Together, these results indicate that loss of Lac1 causes a specific defect in septum organization accompanied by altered cell wall polymers deposition and cytokinesis-associated cell death. The absence of rescue by osmotic stabilization further suggests that the phenotype is not simply attributable to cell wall weakness and increased sensitivity to turgor pressure, but instead it reflects a broader defect in cell envelope organization and morphogenesis.

### Lac1-dependent ceramide synthesis is required for proper localization and activity of β-(1,3)-glucan synthases at the septum

The altered septum ultrastructure and cell wall polymers deposition in *lac1*Δ cells prompted us to ask whether Lac1-dependent ceramide synthesis affects the localization and activity of the cell wall biosynthetic machinery that builds the septum. The *S. pombe* division septum is a distinct three-layered structure comprising a central primary septum flanked symmetrically by secondary septa. Assembly of the primary septum relies on Bgs1/Cps1, which synthesizes linear β-(1,3)-glucan [20,21,50]. To establish proper tripartite septum architecture, Bgs1 operates in concert with other specialized glucan synthases: Bgs4 synthesizes the major branched β-(1,3)-glucan, while Ags1/Mok1 synthesizes α-(1,3)-glucan [15,16,19,51].

We first examined the localization of Bgs4, an essential β-(1,3)-glucan synthase required for cytokinesis and cell wall integrity [51]. In wild-type cells, Bgs4 is localized to active cell growth sites, including the cell tips and division septum. Strikingly, in *lac1*Δ cells, Bgs4 was partially mislocalized, with loss of its normal accumulation at the cell tips in some cells and prominent accumulation in aberrant cytoplasmic granules or vesicles. In contrast, *lag1*Δ cells retained Bgs4 localization at growth sites, although mild intracellular accumulation was also observed (**Figure 2A**).

**Figure 2.**
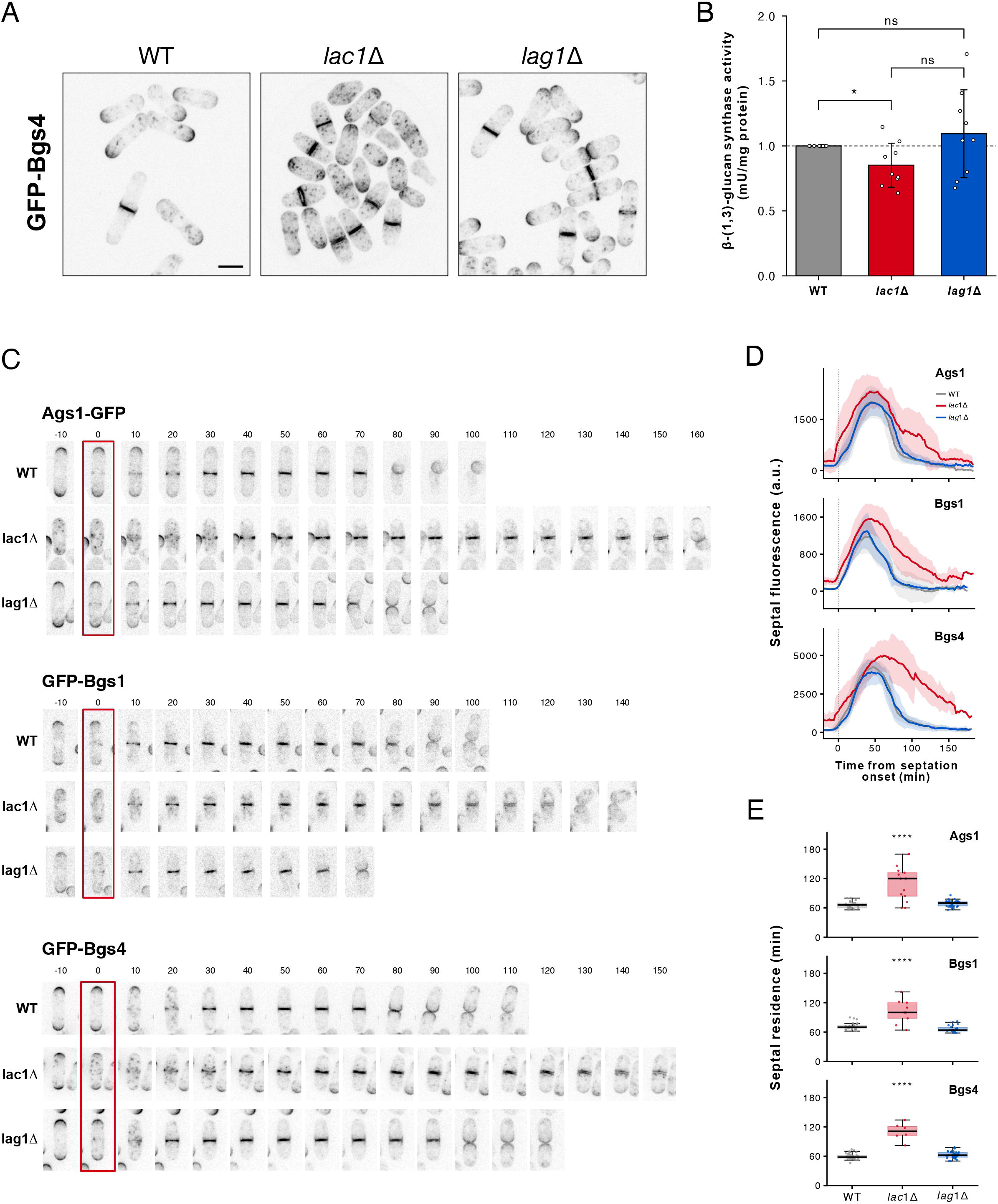
Lac1 is required for the normal localization, activity and septal dynamics of β-(1,3)-glucan synthases. (A) Localization of GFP-Bgs4 in wild-type, *lac1*Δ and *lag1*Δ cells grown at 25°C. Scale bar, 5 µm. **(B)** β-(1,3)-glucan synthase specific activity from membrane extracts, expressed relative to the wild-type specific activity from membrane extract prepared in the same experiment (dashed line, WT = control value of 1). Bars show the mean ± SD of nine independent experiments and points represent individual experiments; wild-type has no associated error by definition of the normalization, as it is the control value of 1 in each experiment. p values are from one-sample t-tests of log-transformed ratios for each mutant against its wild-type control and from a paired t-test between the two mutants; ns, not significant. **(C)** Time-lapse of GFP fluorescence microscopy of Ags1-GFP, GFP-Bgs1 and GFP-Bgs4 in wild-type, *lac1*Δ and *lag1*Δ cells. Images were acquired every 2 min as seven z-sections and are shown as maximum-intensity projections in inverted grayscale. It is shown one frame every 10 min, from 10 min before to 90 min after the onset of septal accumulation (t = 0; see Methods). All nine cells are displayed at the same magnification; grey slots fall beyond the end of the recording. **(D)** Mean septum fluorescence with SD for each synthase oftime-lapses as in C, with every cell aligned on its own onset of septal accumulation; n = 15, 8, and 25 cells (Ags1), 25, 9, and 18 cells (Bgs1), and 28, 10 and 21 cells (Bgs4) for wild-type, *lac1*Δ and *lag1*Δ respectively. a.u.: arbitrary units. **(E)** Elapsed time each synthase remained at the division site, measured from the appearance of septal signal to its loss. Boxes show the median and interquartile range, whiskers the range and circles individual cells; n = 25, 13, and 32 cells (Ags1), 31, 9, and 22 cells (Bgs1) and 38, 6, and 35 cells (Bgs4). Statistical significance was assessed by one-way ANOVA followed by Tukey’s post hoc test; ****, p < 0.0001 for *lac1*Δ against both wild-type and *lag1*Δ. *lag1*Δ did not differ from wild-type for any synthase (p = 0.79, 0.22, and 0.12 for Ags1, Bgs1 and Bgs4, respectively).

Glucan synthases are delivered to the plasma membrane through the secretory pathway [52,53], and ceramides are the precursors of the complex sphingolipids that, together with sterols, organize ordered domains within that membrane. We therefore asked whether mislocalization of Bgs4 in *lac1*Δ cells reflects a general disorganization of sterol-rich membrane domains rather than a defect specific to the glucan synthase machinery. Cells were stained with filipin, a polyene that binds free ergosterol and reports its distribution at the plasma membrane [54]. In wild-type cells, filipin labeled the cell cortex along the whole cell body and was enriched at the growing tips and at the division site. *lag1*Δ cells were indistinguishable from wild-type, and in *lac1*Δ cells filipin also remained confined to the cortex and enriched at the division site, with no accumulation in internal structures, despite oval or rounder cell geometry of this mutant **(Supplementary Figure 3**). The distribution of ergosterol at the plasma membrane is therefore not detectably altered by the loss of Lac1, indicating that the Bgs4 localization defect of *lac1*Δ cells is not a secondary consequence of losing sterol-rich membrane domains. This agrees with previous work showing that the localization of glucan synthases does not depend on sterol-rich domains [55].

To determine whether this altered localization was accompanied by changes in enzymatic output, we measured the *in vitro* β-(1,3)-glucan synthase specific activity in membrane extracts from wild-type, *lac1*Δ and *lag1*Δ cells. Each mutant extract was compared with the wild-type extract prepared in the same experiment, and activities are expressed relative to that control. Across nine independent experiments, activity in *lac1*Δ cells was consistently reduced to 0.85 ± 0.17 of the wild-type value, whereas *lag1*Δ cells showed no consistent change or just a slightly increase (1.09 ± 0.34) (**Figure 2B**). Thus, the partial mislocalization of Bgs4 in *lac1*Δ cells is accompanied by a modest but reproducible reduction in total *in vitro* β-(1,3)-glucan synthase activity, whereas loss of Lag1 affects neither the localization of Bgs4 nor the total *in vitro* activity of the enzyme.

To test whether this defect extends to the septum itself, and to resolve its dynamics, we followed Ags1-GFP, GFP-Bgs1 and GFP-Bgs4 by time-lapse microscopy at 2-min intervals in wild-type, *lac1*Δ and *lag1*Δ cells and quantified their fluorescence at the division site. Each cell was aligned on the onset of septal accumulation (**Figure 2C**). In all three strains every synthase accumulated at the division site, reached a maximum and then declined. In wild-type cells the proteins were cleared as the daughter cells separated, whereas in *lac1*Δ cells they persisted at the division site compared to a wild-type. Quantification of septal fluorescence confirmed this (**Figure 2D**): Bgs1 reached a markedly higher peak in *lac1*Δ cells (+33 % relative to wild-type, against +15 % for Ags1 and +18 % for Bgs4), and all three synthases declined far more slowly after the peak, whereas *lag1*Δ cells were indistinguishable from wild-type. Direct measurement of the time each synthase remained at the septum gave the same result (**Figure 2E**). In wild-type cells Ags1, Bgs1 and Bgs4 persisted for a median of 66, 70 and 58 min respectively, whereas in *lac1*Δ cells all three were significantly prolonged (120, 100 and 111 min, respectively) and much more variable between cells. *lag1*Δ cells resembled wild-type for all three enzymes (70, 64 and 62 min).

Together, these results reveal an apparent paradox: although total *in vitro* β-(1,3)-glucan synthase activity is reduced in *lac1*Δ cell extract, Bgs1 and Bgs4 accumulate to higher levels and persist longer at the septum than in wild-type cells. This argues against a simple model in which septal over-accumulation of glucan synthase proteins reflects increased synthase activity itself. Instead, it suggests that Lac1 loss impairs the turnover or clearance of Bgs1 and Bgs4 from the division site once their local function should be complete. Although the *in vitro* enzymatic activity can be lower, their longer permanence in the membrane can result in a higher accumulation of β-(1,3)-glucan and of secondary septum. This is consistent with the aberrant cytoplasmic accumulation of Bgs4 observed by cell culture imaging (**Figure 2A**), which may reflect mislocalized or improperly recycled enzyme rather than newly synthesized, catalytically active protein.

### Isolation of conditional *lac1* alleles in a *lag1*Δ background

Complete deletion of both *lac1* and *lag1* is inviable under the conditions tested, suggesting that both share a redundant essential function [46,49]. In addition, our previous work showed that the loss of Lac1, but not Lag1, causes substantial reductions in growth rate and cell size. Lipid analysis further suggested that Lac1 and Lag1 contribute differentially to the synthesis of ceramide and complex sphingolipid species [46]. These observations raised the possibility that Lac1 has an essential cellular function that cannot be fully compensated by Lag1.

To investigate the requirement for Lac1-dependent ceramide synthase activity under conditions in which Lac1 function could be acutely perturbed, we performed an unbiased mutagenesis screen to isolate thermosensitive *lac1* alleles in a *lag1*Δ background. We identified multiple *lac1-ts lag1*Δ candidates displaying different degrees of temperature sensitivity when grown on YES plates at 25°C, 28°C, 32°C, and 36°C (**Supplementary Figure 4**). Sequencing of the isolated alleles revealed combinations of single, double, and triple mutations, indicating that multiple residues can influence Lac1 function. Notably, the R37K substitution was present in all of the double-mutant alleles recovered, although these alleles displayed different degrees of temperature sensitivity. We also identified a triple-mutant allele carrying R37K, I143P, and I247L that showed comparatively mild growth defects at the restrictive temperature. These differences in temperature sensitivity, despite the presence of shared mutations, indicate that additional substitutions can modulate the functional consequences of the *lac1* alleles.

The mutations identified in the thermosensitive alleles cluster within the evolutionarily conserved TRAM/LAG1 domain. Alignment of *S. pombe* Lac1 with its *S. cerevisiae* ortholog showed that nearly all of these substitutions fall at residues conserved between the two species, including R37, M167, F175, I247, T255, and I325 (**Supplementary Figure 5**). This conservation, together with their clustering within the TRAM/LAG1 domain, suggests that these residues contribute to a core, evolutionarily conserved function of ceramide synthase. All of the alleles grew comparably to wild-type controls at the permissive temperature (25°C) and therefore provide genetic tools with which to perturb Lac1 function acutely, independently of the chronic adaptation associated with a complete loss of Lac1.

### Acute inactivation of Lac1 reduces ceramide synthesis *in vivo*

A thermosensitive allele is informative only if it compromises the activity of the enzyme it encodes. We therefore asked whether these alleles reduce the output of the ceramide synthase reaction at the restrictive temperature, focusing on *lac1-M167T* and *lac1-R37K L334H*, each alone and in combination with *lag1*Δ deletion. Wild-type and mutant cells were preincubated at 36°C for 1 h and then, labeled with ^[³H]^dihydrosphingosine (DHS) at 36°C for 2 h. Radiolabeled lipids were extracted and separated by thin-layer chromatography (TLC) using solvent system I, which resolves the sphingoid base, ceramide and complex sphingolipid fractions (**Supplementary Figure 6**) [56]. The region of the plate containing ceramides was then scraped, the lipids were re-extracted from the silica gel and resolved in a second TLC step using solvent system II (**Figure 3A**). In our previous work, this second step separated the three fission yeast dihydroceramide species DH-Cer-1, DH-Cer-2 and DH-Cer-3 [46]; because DH-Cer-2 and DH-Cer-3 were not consistently resolved in the present experiment, the three species were quantified together as a single ceramide fraction. For each sample, the signal intensity of the ceramide fraction was divided by the total signal intensity of the second-step TLC, which controls for differences in the uptake of radiolabeled DHS, and the resulting ratio was normalized to the wild-type value of the same replicate.

**Figure 3.**
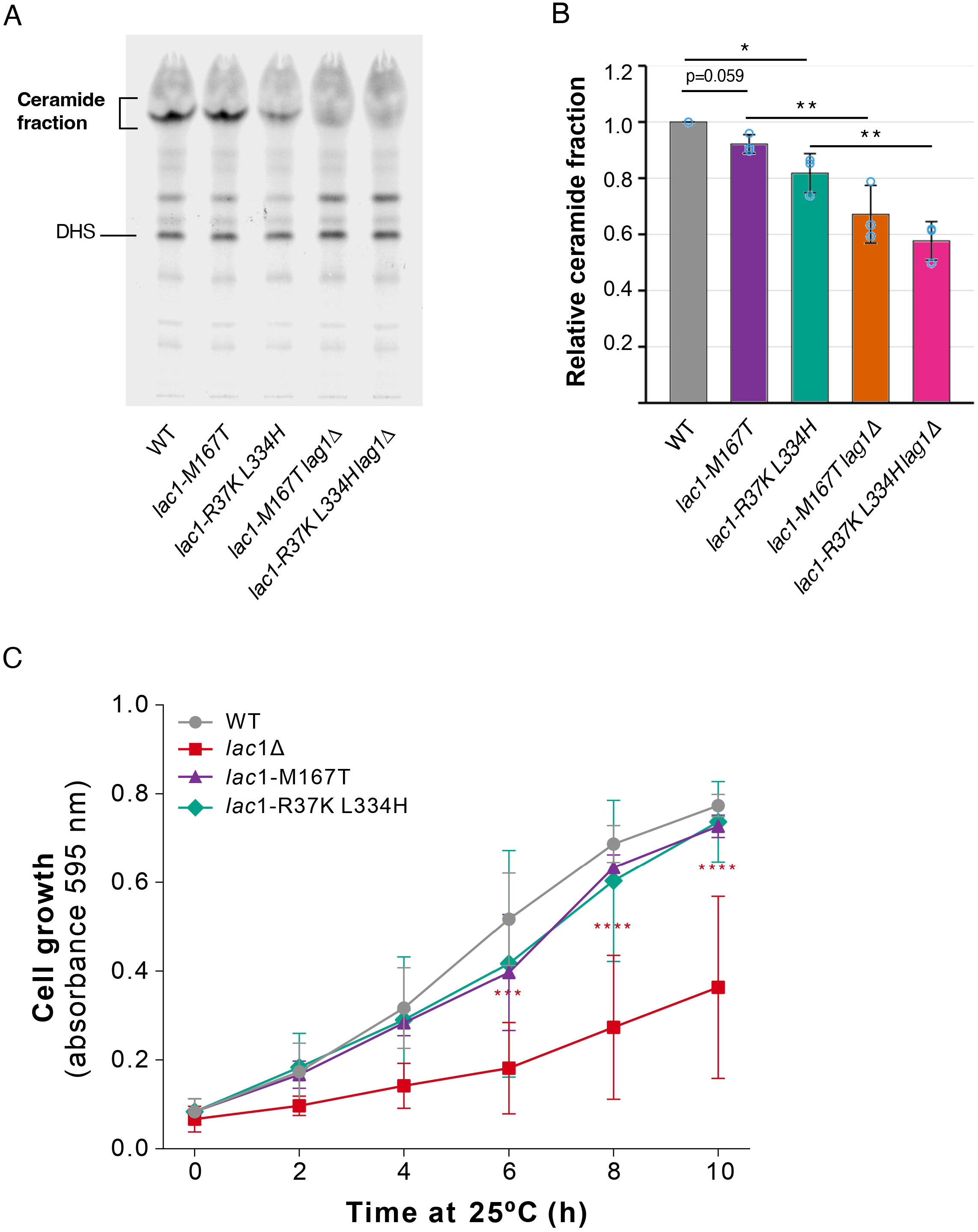
Acute inactivation of Lac1 reduces ceramide synthesis in vivo. **(A)** Representative second-step TLC of the ceramide fraction. Wild-type and the indicated mutant cells were preincubated at 36°C for 1 h and labeled with ^[³H]^DHS at 36°C for 2 h. Radiolabeled lipids were extracted and separated by TLC using solvent system I (**Supplementary** Figure 6); the fractions containing ceramides and dihydrosphingosine (DHS) were collected by scraping the corresponding regions of the plate, eluted from the silica gel with chloroform–methanol, and further separated by TLC using solvent system II. The ceramide fraction indicated includes all ceramide species detected in this assay (DH-Cer-1, DH-Cer-2, and DH-Cer-3; Flor-Parra et al., 2022), which were not consistently resolved as separate bands in this experiment. **(B)** Quantification of the ceramide fraction. For each sample, the signal intensity of the ceramide fraction was divided by the total signal intensity of the second-step TLC, and the resulting ratios were normalized to the corresponding WT value within each replicate, with WT control set to 1. Bars show the mean ± SD of three independent replicates (n = 3); points represent individual replicates. Statistical significance was assessed by one-way ANOVA followed by Tukey’s honestly significant difference test (*, p < 0.05; **, p < 0.01; ns, not significant). **(C)** Cell growth of the indicated strains in liquid YES medium at 36°C, measured as A_595_ over 10 h. Points show the mean of three independent cultures; error bars indicate standard deviation (SD). Statistical significance was assessed by two-way ANOVA with strain and time as factors, followed by Šídák’s test against wild-type at each time point (***, p < 0.001; ****, p < 0.0001); *lac1-M167T* and *lac1-R37K L334H* did not differ from wild-type at any time point.

Acute inactivation of Lac1 reduced the label recovered in the ceramide fraction, and the magnitude of the reduction depended on the presence of Lag1 (**Figure 3B**). In the single mutants, ceramide levels were only modestly affected, reaching 0.92 ± 0.03 of the wild-type value in *lac1-M167T cells* and 0.82 ± 0.07 in *lac1-R37K L334H* cells. Both double mutants with *lag1*Δ, in contrast, showed a pronounced decrease, to 0.67 ± 0.10 in *lac1-M167T lag1*Δ cells and 0.58 ± 0.07 in *lac1-R37K L334H lag1*Δ cells, corresponding to reductions of 33% and 42% relative to wild-type. Each double mutant also differed significantly from the corresponding single mutant, whereas the two double mutants were not significantly different from each other.

We also followed growth in liquid culture at non-permissive temperature. Over 10 h at 36°C, *lac1-M167T* and *lac1-R37K L334H* cells grew with kinetics indistinguishable from wild-type, reaching A_595_ values of 0.73 ± 0.03 and 0.74 ± 0.09 against 0.77 ± 0.03 for wild-type, whereas *lac1Δ* cells reached only 0.36 ± 0.21 **(Figure 3C)**. The conditional substitutions therefore do not compromise growth at 36°C in the presence of Lag1.

These results establish that the *lac1-ts* alleles are hypomorphic for ceramide synthesis at the restrictive temperature and that Lag1 partially compensates for the loss of Lac1 function, which explains why simultaneous deletion of *lag1* is required to uncover their phenotypes (see below). The residual ceramide signal in the double mutants further indicates that the block is partial rather than complete; these strains therefore provide a means of lowering ceramide synthesis acutely rather than of abolishing it.

### Lac1-M167T loses its normal ER localization at the restrictive temperature

Lac1 normally localizes to the endoplasmic reticulum (ER), consistent with its role in ceramide synthesis at this organelle. To determine how the M167T substitution lowers ceramide synthesis, we asked whether it affects the localization of the protein. We examined GFP-tagged Lac1-M167T by fluorescence microscopy at the permissive temperature (25°C) and following a shift to the restrictive temperature (36°C, 4 and 6 h), together with wild-type GFP-Lac1 as a control. In wild-type cells, GFP-Lac1 displayed the expected ER pattern, including perinuclear and peripheral ER staining, at all time points. GFP-Lac1-M167T showed a comparable ER pattern to wild-type at 25°C, but this pattern was progressively lost following the shift to 36°C. Rather than the continuous perinuclear and peripheral ER network seen in wild-type cells, GFP-lac1-M167T accumulated in discrete, vesicle-like puncta clustered around the nucleus (**Figure 4A**).

**Figure 4.**
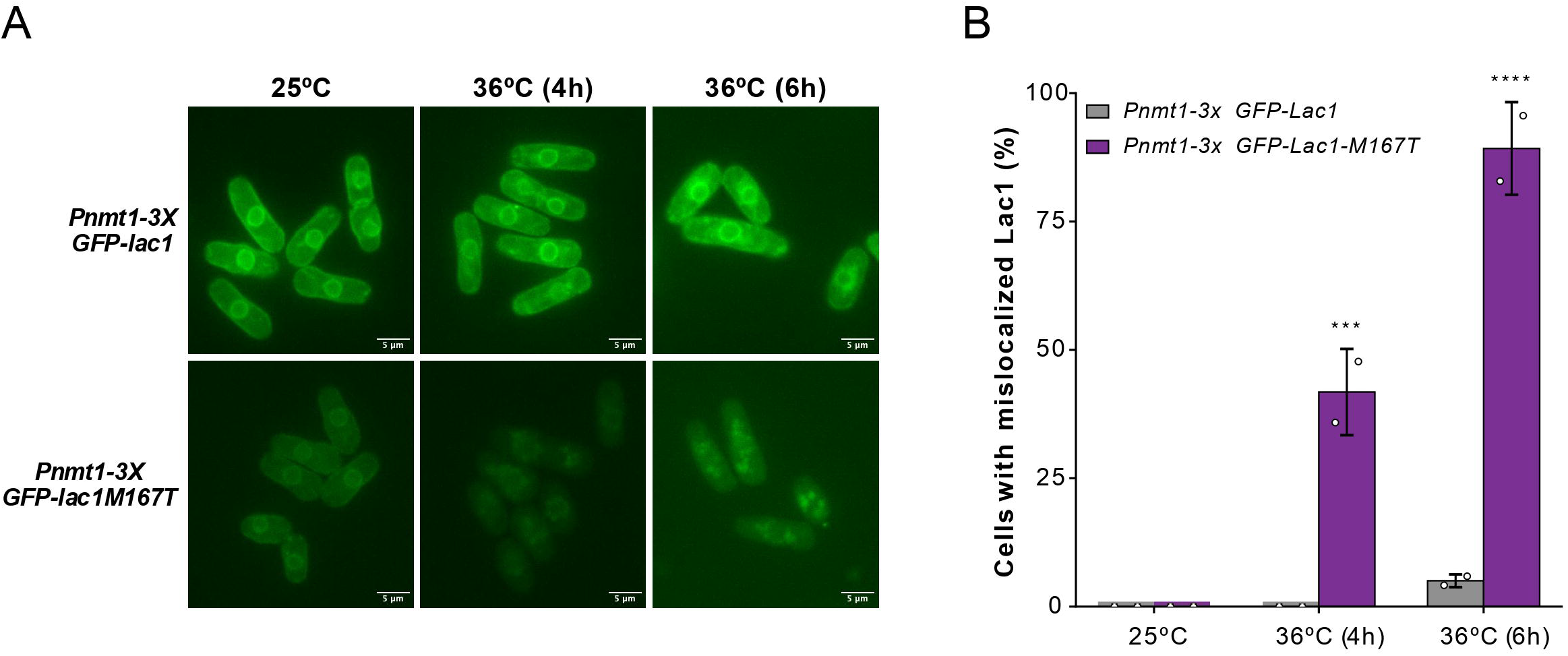
The lac1-M167T protein loses its endoplasmic reticulum localization at the restrictive temperature. **(A)** Localization of wild-type GFP-Lac1 and thermosensitive GFP-Lac1-M167T in cells grown at 25°C and after 4 and 6 h at 36°C. Both proteins are expressed from the 3X version of the *nmt1^+^* promoter (*nmt1-3X*). Scale bar, 5 µm. **(B)** Percentage of cells showing a mislocalized, punctate GFP-Lac1 pattern at each time point. Bars show the mean ± SD of two independent experiments with 400 cells scored per condition in each case; circles show the individual experiments. Statistical significance was assessed by ordinary two-way ANOVA with genotype and time as factors (interaction, genotype and time each p < 0.0001), followed by Bonferroni’s multiple comparisons test between genotypes within each time point (***, p < 0.001; ****, p < 0.0001); the two strains did not differ at 25°C (p > 0.9999).

Quantification of this phenotype confirmed a significant, temperature-dependent increase in Lac1 mislocalization specific to the M167T allele. Wild-type GFP-Lac1 showed essentially no mislocalization at 25°C or after 4 h at 36°C, with only a small fraction of cells (5.0 %) showing a mislocalized pattern after 6 h at 36°C. In contrast, GFP-Lac1-M167T showed no mislocalization at 25°C, but the proportion of cells displaying a mislocalized, punctate pattern rose sharply to 41.8% after 4 h at 36°C and to 89.2% after 6 h at 36°C (**Figure 4B**). The M167T substitution therefore compromises the ability of Lac1 to maintain its normal ER localization at elevated temperature, providing a likely mechanistic basis for the reduced ceramide synthesis measured in this allele.

### Structural modeling suggests that the two alleles perturb distinct contacts with acyl-CoA

The two conditional alleles R37K L334H and M167T, reduced ceramide synthesis to a similar extent, yet they affect residues that lie far apart in the TRAM/LAG1 domain. To ask whether they compromise the enzyme in the same way, we built a structural model of the *S. pombe* Lac1-C_26_-CoA complex. The AlphaFold model of SpLac1 was superimposed on the Lac1 subunit of the C_26_-CoA-bound *S. cerevisiae* Lac1-Lip1 complex (PDB 8IZD; [57]), the experimentally observed C_26_-CoA coordinates were transferred to the SpLac1 model, and residues 58–353 of the 384-residue protein, corresponding to the high-confidence core, were embedded in a pre-equilibrated yeast ER membrane (**Figure 5A, B**). Because no Lip1 ortholog has been identified in *S. pombe*, only the Lac1 subunit was simulated. Three independent 200-ns molecular dynamics (MD) simulations were performed per genotype at 36°C, the temperature at which the conditional alleles lose function.

**Figure 5.**
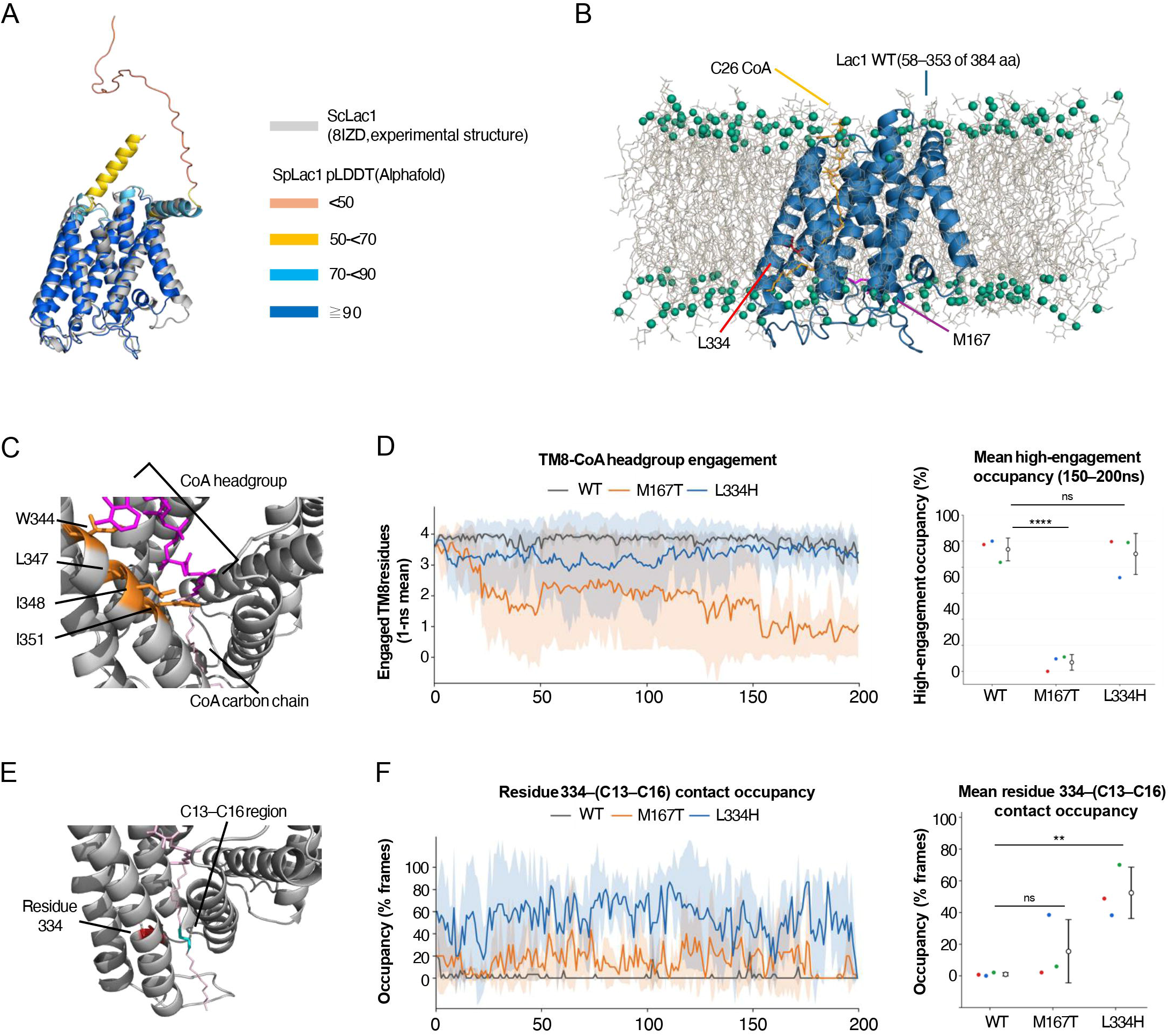
Structural modeling and molecular dynamics analysis of wild-type and mutant SpLac1–C26-CoA complexes. **(A)** Superposition of the AlphaFold model of *Schizosaccharomyces pombe* Lac1 (SpLac1; UniProt O59735), colored by predicted local distance difference test (pLDDT) score as indicated, and the Lac1 subunit of the experimental cryo-EM structure of the *Saccharomyces cerevisiae* Lac1–Lip1 complex (ScLac1; PDB 8IZD, gray). **(B)** Membrane-embedded wild-type (WT) SpLac1–C_26_-CoA model used for molecular dynamics (MD) simulations. Lac1 is shown as a blue cartoon and comprises residues 58–353 of the full-length 384-residue protein; no Lip1 ortholog has been identified in *S. pombe* and none was included in the simulated system. C_26_-CoA, M167 and L334 are highlighted in yellow, purple, and red, respectively. Membrane lipids of the pre-equilibrated yeast ER bilayer are shown as gray lines, with their headgroups represented by green spheres. **(C)** WT TM8 region, with W344, L347, I348, and I351 highlighted in orange. The CoA headgroup and acyl chain of C_26_-CoA are shown in magenta and pink, respectively. **(D)** TM8–CoA headgroup engagement. Left, time course of the number of engaged residues among W344, L347, I348, and I351 over 0–200 ns, averaged in nonoverlapping 1-ns bins. A residue was considered engaged when its minimum heavy-atom distance to the CoA headgroup was <0.45 nm. Right, high-engagement occupancy over 150–200 ns, defined as the percentage of time with three or four of these residues simultaneously engaged. n.s= nor significant; ** p = 0.0078 **(E)** Close-up view of WT residue L334 (red) and C_26_-CoA (pink), with the C_13_–C_16_ acyl-chain segment highlighted in cyan. Acyl carbons are numbered from the thioester carbonyl carbon, C1. **(F)** Contact occupancy between residue 334 (Leu in WT and M167T; His in L334H) and acyl carbons C_13_–C_16_ of C_26_-CoA. A contact was defined by a minimum distance <0.45 nm between these carbons and residue 334 heavy atoms, including backbone atoms. Occupancy was calculated as the percentage of frames containing a contact. Left, occupancy in nonoverlapping 1-ns bins over 0–200 ns; right, occupancy over the full 0–200-ns interval. Three independent 200-ns MD simulations were analyzed per genotype (n = 3). n.s= nor significant; **** p = 0.0001. In the time-course plots in **(D)** and **(F)**, lines and shaded regions indicate the mean and SD across trajectories, respectively; WT, M167T, and L334H are shown in gray, orange, and blue, respectively. In the summary plots in **(D)** and **(F)**, red, blue, and green points represent individual trajectories, and black open circles with error bars indicate the mean ± SD. Genotypes were compared separately for each summary outcome using one-way ANOVA followed by Tukey’s honestly significant difference test. Tukey-adjusted P values for comparisons with WT are shown.

Neither mutation significantly altered the mean backbone RMSD of the simulated Lac1 protein, the modeled portion of the TRAM/LAG1 domain, or individual transmembrane helices (TM1–TM8) over 0–200 ns (**Supplemental Figure 7A**). We then examined the catalytic region. Neither mutation altered the local backbone conformation of the H228, H229, D256 and D259 cluster, the distance between H228 Nε2 and the thioester carbonyl carbon (C1) of C_26_-CoA, or the occupancy of that proximity (**Supplemental Figure 7B, C**). The mutants instead differed in how they contacted the substrate, and they did so at different positions. M167T reduced the fraction of time during which at least three of the four TM8 residues W344, L347, I348 and I351 simultaneously contacted the CoA headgroup (p = 0.0001 versus wild-type over the 150–200 ns window), whereas L334H was indistinguishable from wild-type in this measure (p = 0.9205) (**Figure 5C, D**). Conversely, L334H increased the contact occupancy between residue 334 and the C13–C16 segment of the acyl chain (p = 0.0078 versus wild-type), whereas M167T did not (**Figure 5E, F**). Because M167 lies outside TM8, its effect on TM8-CoA engagement is necessarily indirect. These simulations therefore identify two distinct perturbations of the Lac1-substrate interface, one at the CoA headgroup and one at the acyl chain, rather than a shared defect in the catalytic center.

### *lac1-ts* alleles cause temperature-dependent cell death in the absence of Lag1

We next asked how the acute reduction in ceramide synthesis affects cell viability. We performed serial dilution spot assays on plates containing phloxine B, a vital dye that accumulates in dead or dying cells, allowing growth and viability to be assessed simultaneously, across a temperature range from permissive (25°C) to restrictive (36°C) (**Figure 6A**).

**Figure 6.**
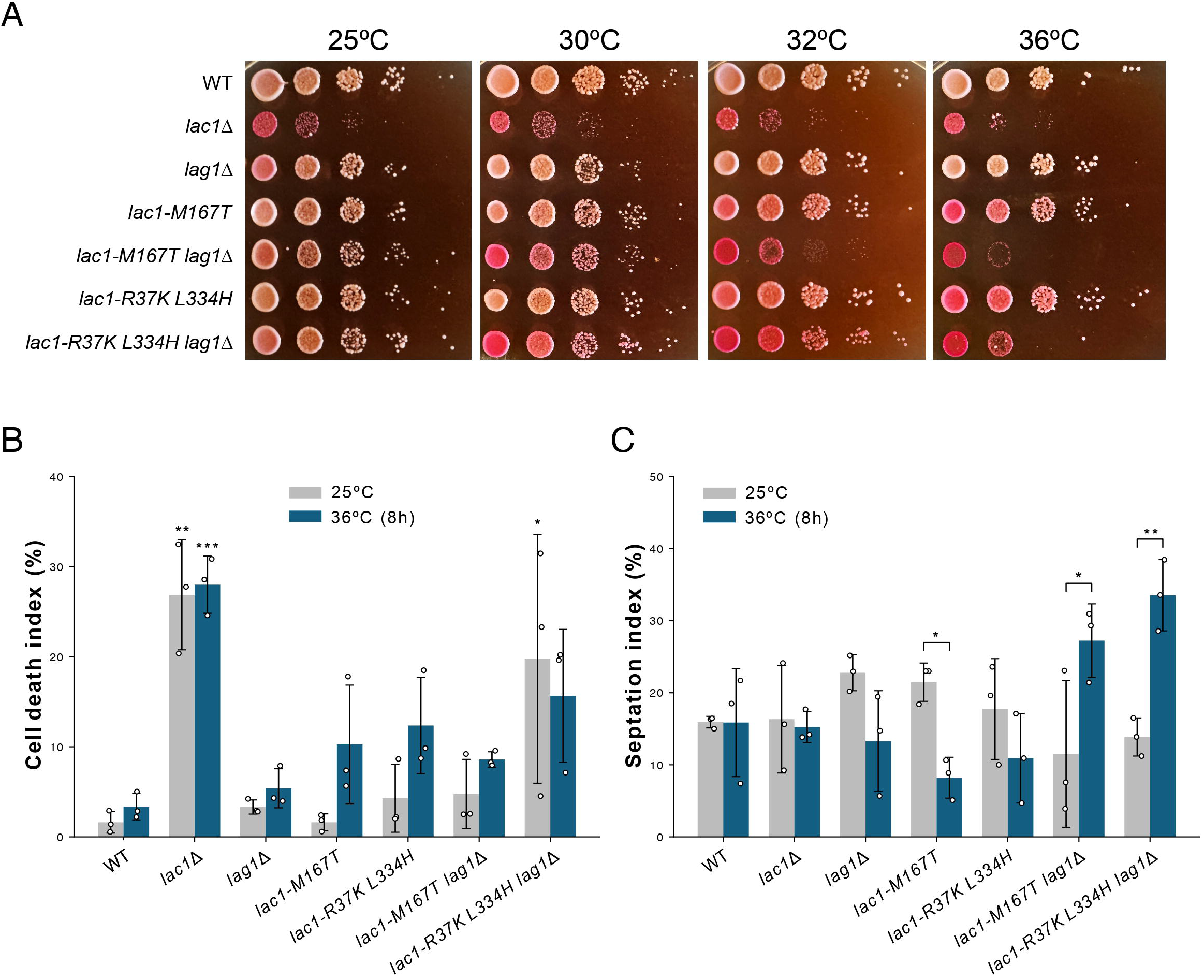
Conditional loss of Lac1 causes cell death and a septation defect at the restrictive temperature. **(A)** Serial-dilution spot assay of the indicated strains. Ten-fold dilutions were spotted onto YES medium containing phloxine B and incubated at 25°C, 30°C, 32°C and 36°C for 2-4 days. Phloxine B accumulates in dead cells, staining the colony to pink color. **(B)** Cell death index, scored as the percentage of cells that showed intense calcofluor staining in cells grown at 25°C and after 8 h at 36°C. **(C)** Septation index, scored as the percentage of cells with a septum, in the same cultures as in B. In (B) and (C) bars show the mean ± SD of three independent experiments and circles show the individual experiments (n> 400 cells/experiment). Because each experiment measured both temperatures, data were analyzed by two-way ANOVA with strain as a between-experiment factor and temperature as a repeated measure, followed by Šídák’s test comparing 25°C with 36°C within each strain. For the cell death index, strain p = 0.0001, temperature p = 0.017 and the strain × temperature interaction p = 0.11; no within-strain comparison between temperatures reached significance after correction for the seven strains. For the septation index, the strain × temperature interaction p < 0.0001, and the septation index differed between temperatures in *lac1-M167T* (p = 0.015), *lac1-M167T lag1*Δ (p = 0.004) and *lac1-R37K L334H lag1*Δ (p = 0.0005). Comparisons without an asterisk are not significant. * p ≤ 0.05, ** p ≤ 0.01, *** p ≤ 0.001.

At 25°C, wild-type, *lag1*Δ, both single mutants (*lac1-M167T* and *lac1-R37K L334H*), and both double mutants (*lac1-M167T lag1*Δ and *lac1-R37K L334H lag1*Δ) all grew normally with minimal phloxine B staining. *lac1*Δ cells were the only strain to show the characteristic increase in pink staining and reduced colony number consistent with the elevated cell death reported above (**Figure 1A**). On plates, therefore, *lac1*Δ was the only strain affected at this temperature, although the quantitative death index showed that *lac1-R37K L334H lag1*Δ cells already die at an appreciable rate at 25°C (**Figure 6B**). At 30°C, *lag1*Δ and the single mutants remained unaffected, whereas both double mutants already showed a visible increase in phloxine B staining, despite still forming colonies across the full dilution series comparable to wild-type. Across the full temperature range tested, *lag1*Δ cells remained unaffected, with minimal phloxine B staining and colony formation comparable to wild-type at every temperature. The *lac1-M167T* and *lac1-R37K L334H* single mutants began to show increased phloxine B staining at 32°C, though still less intense than *lac1*Δ and the double mutants at this temperature, and reached staining levels comparable to *lac1*Δ by 36°C; despite this increased staining, both single mutants continued to form colonies similar to wild-type at all temperatures, indicating that increased cell death did not translate into a cell growth defect. The double mutants, in contrast, progressively diverged from wild-type: at 32°C, *lac1-M167T lag1*Δ cells already resembled *lac1*Δ in both phloxine B staining and reduced colony number, whereas *lac1-R37K L334H lag1*Δ cells showed phloxine B staining comparable to *lac1*Δ but still formed colonies similar to wild-type at this temperature. At 36°C, both double mutants were indistinguishable from *lac1*Δ in phloxine B staining and loss of colony growth.

These results show that the reduction in ceramide synthesis caused by the *lac1-ts* alleles translates into cell death rather than simple growth arrest, and that this outcome requires the simultaneous loss of Lag1. The two alleles differ in which assay reports the more severe defect: *lac1-M167T lag1*Δ shows the stronger colony-level phenotype, becoming indistinguishable from *lac1*Δ already at 32°C, whereas *lac1-R37K L334H lag1*Δ shows the higher cell death and septation index (**Figure 6B, C**), and the lower ceramide signal (**Figure 3B**). Notably, *lac1-R37K L334H* single-mutant cells showed a measurable reduction in ceramide synthesis (**Figure 3B**), yet formed colonies indistinguishable from wild-type at all temperatures, indicating that fission yeast tolerates a moderate decrease in ceramide production, and that viability is compromised only when synthesis falls below a threshold.

To quantify this temperature-dependent death directly, we measured the cell death index of all seven strains at 25°C and after 8 h at 36°C (**Figure 6B**). At 25°C, cell death was low in wild-type (1.6%), *lag1*Δ (3.3%), *lac1-M167T* (1.6%), lac1*-R37K L334H* (4.3%) and *lac1-M167T lag1*Δ (4.8%) cells, while *lac1*Δ cells showed the expected elevated death index (26.9 ± 6.1%), consistent with our earlier results (**Figure 1A**). *lac1-R37K L334H lag1*Δ cells were the exception among the conditional strains, reaching 19.3 ± 10.2% already at the permissive temperature, although with considerable variation between experiments. Following the shift to 36°C for 8 h, cell death tended to increase in both single thermosensitive mutants (*lac1-M167T*: 10.3%; *lac1-R37K L334H*: 12.4%) and double mutants (*lac1-M167T lag1*Δ: 8.6%; *lac1-R37K L334H lag1*Δ: 15.6%), although none of these increases reached significance, while wild-type and *lag1*Δ cells remained low (3.4% and 5.4%, respectively) and *lac1*Δ cells remained high (28.0%). Notably, the magnitude of death measured at 8 h was more modest than the severe colony-level growth defects seen on plates, particularly for *lac1-M167T lag1*Δ, which showed the most severe plate phenotype (**Figure 6A**) despite a comparatively modest cell death index at this timepoint. This raises the possibility that the growth arrest in this allele reflects cell-cycle arrest in addition to, or preceding, cell death, a hypothesis we have not yet directly tested.

### Acute loss of Lac1 function causes septation defects

To test whether this temperature-dependent death is likewise associated with a failure to complete cell division, as in *lac1*Δ cells, which die during cell separation and assemble septa with an abnormal layered architecture (**Figure 1**), we measured the septation index of all seven strains at 25°C and after 8 h at 36°C (**Figure 6C**). An increased septation index indicates that cells accumulate in the septated state, either because septum assembly and maturation are slowed or because cell separation is delayed. At 25°C, septation indexes were broadly similar across strains, ranging from 11.5% (*lac1-M167T lag1*Δ) to 22.8% (*lag1*Δ), with wild-type at 15.9% and *lac1Δ* at 16.3%. Following the shift to 36°C for 8 h, the septation index did not increase in wild-type, *lag1*Δ, or either single thermosensitive mutant (wild-type: 15.9%; *lag1*Δ: 13.3%; *lac1-M167T*: 8.2%; *lac1-R37K L334H*: 10.9%), nor *lac1*Δ cells (15.2%, similar to its own 25°C value); in *lac1-M167T* cells the index in fact decreased significantly after the shift (21.5% to 8.2%). In contrast, both double mutants showed a marked increase in septation index: *lac1-M167T lag1*Δ rose from 11.5% to 27.2%, and *lac1-R37K L334H lag1*Δ rose from 13.9% to 33.5%, the highest of any strain or condition tested. This selective accumulation of septated cells specifically in the double mutants at the restrictive temperature, indicates that acute loss of Lac1 function causes a cytokinesis defect, consistent with the cell death occurring preferentially during or after septum degradation, and with the defects in septum organization and cell wall polymers deposition observed in cells chronically deprived of Lac1 (**Figure 1**).

## DISCUSSION

How fungal cells couple the expansion of the plasma membrane to the synthesis and remodeling of the cell wall remains one of the central unresolved questions in fungal morphogenesis [3,5]. The results presented here identify ceramide synthesis as one of the inputs to that coupling. Cells deprived of Lac1 die predominantly during cell separation, assemble division septa with an abnormal layered architecture, deposit altered amounts of cell wall polymers, and fail to clear the glucan synthases that build the septum once division is complete. Acute inactivation of Lac1 reproduces the cytokinesis-associated death and the accumulation of septated cells, and we show directly that these conditional alleles lower ceramide synthesis at the restrictive temperature. Together, these observations argue that Lac1-dependent ceramide synthesis is required for the orderly assembly and completion of the division septum, in addition to the cell wall synthesis as such.

The most informative aspect of the *lac1*Δ phenotype is its internal contradiction. Total *in vitro* β-(1,3)-glucan synthase activity measured in membrane extracts is reduced, and Bgs4 is partly diverted into cytoplasmic granules, yet Ags1, Bgs1 and Bgs4 reach higher peak levels at the division site and remain there roughly twice as long as in wild-type cells. Over-accumulation of these enzymes at the septum is therefore not a consequence of increased synthetic capacity. The simplest interpretation is that loss of Lac1 impairs the removal of glucan synthases from the division site once their local task is complete, so that catalytically compromised or improperly recycled enzyme persists where it should have been cleared. Such a defect would account both for the thicker secondary septa seen by electron microscopy and for the increased aniline blue staining, and it predicts that the primary lesion lies in the membrane environment through which these enzymes are delivered and retrieved rather than in their catalytic chemistry. A similar defect of lower *in vitro* β-(1,3)-glucan synthase activity while increasing and retaining the Ags1, Bgs1, and Bgs4 localization in the plasma membrane, thus causing a long-term β-(1,3)-glucan and cell wall accumulation, has been described in cells treated with the antifungal theonellamide, whose primary target are the plasma membrane sterols [58]. Because the distribution of ergosterol is unchanged in *lac1Δ* cells, this environment cannot be equated with sterol-rich domains, in agreement with the finding that glucan synthase localization does not depend on them [55]. Ceramides and the complex sphingolipids derived from them are instead candidates for a distinct, sterol-independent requirement, consistent with the role of very-long-chain ceramides in the sorting and transport of membrane proteins [37].

A second question raised by our previous work was why the two catalytic subunits of the same enzyme are not functionally equivalent [46]. Cells lacking Lag1 were indistinguishable from wild-type in the frequency of death during cell separation, in septum ultrastructure, in aniline blue, calcofluor white and FITC-lectin staining of cell wall polymers, in the distribution of plasma membrane sterols, and in total *in vitro* β-(1,3)-glucan synthase activity; the only differences we detected were a mild intracellular accumulation of Bgs4 and a slightly shorter residence of Bgs1 at the division site. This is not the consequence of an unperturbed pathway, since *lag1*Δ cells do not synthesize the long-chain dihydroceramides DH-Cer-2 and DH-Cer-3, whereas *lac1*Δ cells retain both and instead fail to produce phytoceramide [46]. Two strains with substantially altered but different ceramide profiles therefore diverge completely in phenotype, which argues that the cell wall and cytokinesis defects described here do not follow from a general reduction in ceramide output but from the loss of the specific products of Lac1. The metabolic labeling presented here offers a complementary answer at the level of total pathway flux: Lag1 can sustain a substantial fraction of ceramide synthesis when Lac1 is inactivated, and the strongest reductions are obtained only when both subunits are compromised. At the same time, ceramide synthesis is not simply additive with viability. Cells carrying a single conditional allele show a measurable decrease in labeled ceramide, yet forming colonies indistinguishable from wild-type, whereas the double mutants, in which synthesis falls by a third to nearly one half, arrest and die. Fission yeast therefore tolerates a considerable reduction in ceramide production, and the cell wall and cytokinesis phenotypes appear only once synthesis falls below a threshold. This threshold behavior also explains why the complete loss of Lac1 is tolerated at all: Lag1 maintains enough flux for growth, but not enough to support faithful septum assembly, which is precisely the process that fails in *lac1*Δ cells.

Comparison of the chronic and acute perturbations is instructive in both directions. Cells that have lived without Lac1 for many generations may have remodeled their lipid composition and their cell wall, so that part of the *lac1*Δ phenotype could reflect adaptation rather than the immediate consequence of ceramide loss. The conditional alleles bypass this adaptation, and the phenotypes they produce at the restrictive temperature (growth arrest, septated cells and death) match the deletion phenotype. Conversely, the acute perturbation is partial and short, which may explain why the cell death index measured after 8 h is more modest than the colony-level growth defect on plates. The possibility that acute Lac1 inactivation first imposes a cell-cycle arrest, with death following later, remains to be tested in the future.

### Lac1 mutant function and substrate positioning

The two conditional alleles characterized here reduce ceramide synthesis to a comparable degree, and the MD simulations add a structural interpretation of how they may do so by identifying changes in contact with different parts of C_26_-CoA. Ceramide synthesis transfers the acyl group of acyl-CoA to a sphingoid base and releases CoA-SH [57], so both the accommodation of the acyl chain and the positioning of the CoA moiety can contribute to maintaining a substrate configuration compatible with acyl transfer.

M167T reduced the fraction of time during which at least three of the four examined TM8 residues simultaneously contacted the CoA headgroup, which suggests a change in the contact network that positions the CoA portion of the substrate; reduced coordinated contacts with the headgroup could make productive orientations of acyl-CoA less frequent, even though CoA is released during the reaction. L334H showed no significant difference from wild-type in this measure, and instead increased contact between residue 334 and the central C_13_–C_16_ segment of the acyl chain, identifying the local environment around the acyl chain as a candidate contributor to this allele. Together with its unchanged catalytic-region readouts, this suggests that the Leu-to-His substitution may change chain packing and make substrate positioning less favorable for the reaction. The selected catalytic-region readouts showed no significant wild-type– mutant differences, so the clearest structural signatures detected here concern substrate contacts rather than the catalytic center itself.

These results support a model in which the two substitutions converge on a similar reduction in ceramide output through distinct changes at the Lac1-substrate interface: M167T may impair positioning of the CoA headgroup, while L334H may alter accommodation of the acyl group that is transferred into ceramide. That two residues on opposite sides of the substrate-binding environment can be perturbed to similar functional effect is consistent with their clustering in the conserved TRAM/LAG1 domain and brings the organization of the substrate-binding environment into the discussion of Lac1 function.

These interpretations carry the limitations inherent to the approach. The simulations model the L334H substitution in isolation, whereas the corresponding strain also carries R37K; residue 37 lies outside the modeled construct (residues 58–353), so its contribution was not assessed. Only the Lac1 subunit was simulated: the template structure is a Lac1-Lip1 complex, and although no Lip1 ortholog has been identified in *S. pombe*, a contribution of an accessory subunit to the substrate environment cannot be excluded. The analysis rests on three 200-ns trajectories per genotype and on contact-geometry readouts, with C_26_-CoA parameters extrapolated from a C_22_-CoA model and a neutral HSD tautomer assumed for the introduced H334. Proposed effects on productive substrate positioning are therefore mechanistic interpretations of contact geometry rather than measurements of reaction rate.

### Concluding remarks

Our data place ceramide synthesis upstream of the spatial control of cell wall assembly and identify the clearance of glucan synthases from the division site as the step most clearly affected. Several questions follow. It remains to be established which lipid species carries the requirement, since ceramides are both bioactive lipids in their own right and the precursors of inositol-phosphoceramide and its mannosylated derivative, and our previous lipid analysis showed that the loss of Lac1 alters the whole downstream profile [46]. It also remains open whether the persistence of Ags1, Bgs1 and Bgs4 at the septum reflects defective endocytic retrieval, altered interaction with the exocyst and the septation machinery, or a change in the physical properties of the plasma membrane at the division site. Because the cell wall is essential for fungal viability and absent from mammalian cells, and because sphingolipid metabolism is itself a validated antifungal target, the pathway described here may offer an additional point of intervention [26]. Finally, ceramide synthase subunits are conserved from yeast to humans, and the residues mutated in our conditional alleles lie in the conserved TRAM/LAG1 domain, so the substrate-positioning defects suggested by our simulations may be relevant beyond fission yeast.

**Supplementary Figure 1. Osmotic stabilization does not rescue the cell growth defect of *lac1*Δ cells.** Ten-fold serial dilutions of the indicated strains were spotted onto YES and YES supplemented with 1.2 M sorbitol and incubated at the indicated temperatures for 2-3 days.

**Supplementary Figure 2. Loss of Lac1 causes altered septum wall polymers deposition as observed by fluorochrome staining.** Septum wall polymers staining. The fluorescence intensities of cells as in figure 1C, 1D, and 1E, were analyzed by Linescan along the entire septum. Left panels: *lac1*Δ, but not *lag1*Δ, cells show higher aniline blue (AB) fluorescence along the septum compared to that of wild type cells. Right panels: *lac1*Δ, *lag1*Δ, and wild-type cells present similar calcofluor white (CW) fluorescence along the septum. Quantification of mean fluorescence intensity in a line along the septum compared to the wild-type (WT) (number of cells: Left, AB: *lac1*Δ/WT-RFP: 22/15; *lag1*Δ/WT-RFP: 8/8; WT/WT-RFP: 10/10. Right, CW: *lac1*Δ/WT-RFP: 5/6; *lag1*Δ/WT-RFP: 8/10; WT/WT-RFP: 10/5). Bars show the mean ± SD. a.u.: arbitrary units

**Supplementary Figure 3. Loss of Lac1 does not detectably alter the distribution of plasma membrane sterols.** Wild-type, *lac1*Δ and *lag1*Δ cells were grown in YES medium at 25°C, stained with filipin and imaged by fluorescence microscopy. In all three strains, filipin labels the cell cortex along the cell body and is enriched at the growing tips and at the division site, with no accumulation in internal structures; *lac1*Δ cells display the oval and rounder geometry characteristic of this mutant. The uniformly bright cell in the *lac1*Δ field is a dead or dying cell. Images are single focal planes from one representative experiment of three. Scale bar, 5 µm.

**Supplementary Figure 4. Isolation of thermosensitive lac1 alleles in a *lag1*Δ background.** Ten-fold serial dilutions of wild-type and of the *lac1-ts lag1*Δ candidates recovered from the mutagenesis screen were spotted onto YES plates and incubated at 25°C, 28°C, 32°C and 36°C for 2-4 days.

**Supplementary Figure 5. The thermosensitive Lac1 substitutions affect residues within the conserved TLC domain of ceramide synthase.** Multiple sequence alignment of *Schizosaccharomyces pombe* Lac1 with ceramide synthase orthologs from *Homo sapiens* (CERS2, UniProt Q96G23), *Mus musculus* (Cers2, Q924Z4), *Drosophila melanogaster* (schlank, Q9W423), *Caenorhabditis elegans* (hyl-1, G5ED45), and *Saccharomyces cerevisiae* (Lac1, P28496); *S. pombe* Lac1 is O59735. CERS2 was chosen as the vertebrate representative because, like Lac1, it uses very-long-chain acyl-CoAs. Full-length sequences were aligned with MAFFT v7 using the --auto strategy (Katoh and Standley, 2013), and the region corresponding to residues 160–367 of *S. pombe* Lac1 is shown in five blocks; the numbers flanking each row give the first and last residue of that sequence in the block. Residues mutated in the thermosensitive alleles are boxed in blue and labeled in red above the alignment, with the substitution and the position given in *S. pombe* numbering. The catalytic histidine pair (H228, H229) and the aspartate pair (D256, D259) analyzed by molecular dynamics (**Figure 5**) are boxed in purple and labeled in orange. Below each block, red bars mark the transmembrane helices annotated for *S. pombe* Lac1 in UniProt (TM3–TM7 of seven; TM1 and TM2 lie N-terminal to the displayed region) and the green bar marks the TLC/TRAM-LAG1-CLN8 domain (InterPro IPR006634), which spans residues 141–357. The R37K substitution is not shown because residue 37 lies N-terminal to the displayed region, within a predicted disordered segment (residues 19–42) outside the TLC domain. Relative to *S. cerevisiae* Lac1, *S. pombe* Lac1 is identical at M167, F175, F244, T254, and I325, as well as at the catalytic H228, H229, D256, and D259, and carries conservative substitutions at I247 (Met in ScLac1), M290 (Leu), L334 (Ile), and T335 (Gly).

**Supplementary Figure 6. Separation of sphingolipid species after ^[³H]^DHS labeling.** Wild-type and mutant cells were preincubated at 36°C for 1 h and then, labeled with ^[³H]^dihydrosphingosine (DHS) at 36°C for 2 h. Radiolabeled lipids were extracted and separated by thin-layer chromatography (TLC) using solvent system I. Dashed boxes indicate the fractions containing ceramides and DHS.

**Supplemental Figure 7. Backbone RMSD and catalytic-region geometry of wild-type and mutant SpLac1–C_26_-CoA complexes. (A)** Mean backbone root-mean-square deviation (RMSD) over 0–200 ns for the simulated Lac1 protein (Whole; residues 58–353), the modeled portion of the TRAM/LAG1 domain (residues 141–353), and TM1–TM8, with the residue ranges indicated. RMSD was calculated for the backbone N, CA, and C atoms of each region after equally weighted least-squares fitting of the backbone atoms of residues 58–353 to the first production frame of each trajectory. The same whole-protein fit was used for all regions. **(B)** WT catalytic region, with H228 and H229 highlighted in orange, D256 in blue, and D259 in turquoise. C_26_-CoA is shown as pink sticks, with its thioester carbonyl carbon (C1) represented as a sphere. The red dashed line indicates the H228 Nε2–C1 distance. **(C)** Comparison of WT, M167T, and L334H over 0–200 ns. Left, mean catalytic-core local backbone root-mean-square deviation (RMSD) of H228, H229, D256, and D259, calculated after fitting the same backbone atoms to the first production frame of each trajectory. Middle, mean distance between H228 Nε2 and the C_26_-CoA thioester carbonyl carbon (C1). Right, H228 Nε2–C1 proximity occupancy, defined as the percentage of frames with a distance ≤0.45 nm. Three independent 200-ns MD simulations were analyzed per genotype (n = 3). In (A), WT, M167T, and L334H are shown in red, blue, and green, respectively, and colored points represent individual trajectory means. In (C), red, blue, and green points represent individual trajectories. Black open circles with error bars in (A) and (C) indicate the mean ± SD across trajectories. Genotypes were compared separately for each region in (A), and each summary outcome in (C), using one-way ANOVA followed by Tukey’s honestly significant difference test. Tukey-adjusted P values for comparisons with WT are shown.

## MATERIALS AND METHODS

### Yeast strains and media

All yeast strains used in this study are listed in Table 1. Cells were grown at 25°C or 30°C in standard YES media (MPBiomedical, Irvine, CA, USA), supplemented with 200 mg/L adenine, histidine, leucine, uracil, and lysine (Sigma-Aldrich, St. Louis, MO, USA). Plates were made by adding 2% agar (Condalab). Standard methods for *S. pombe* were used for strain constructions [59].

**Table 1.**
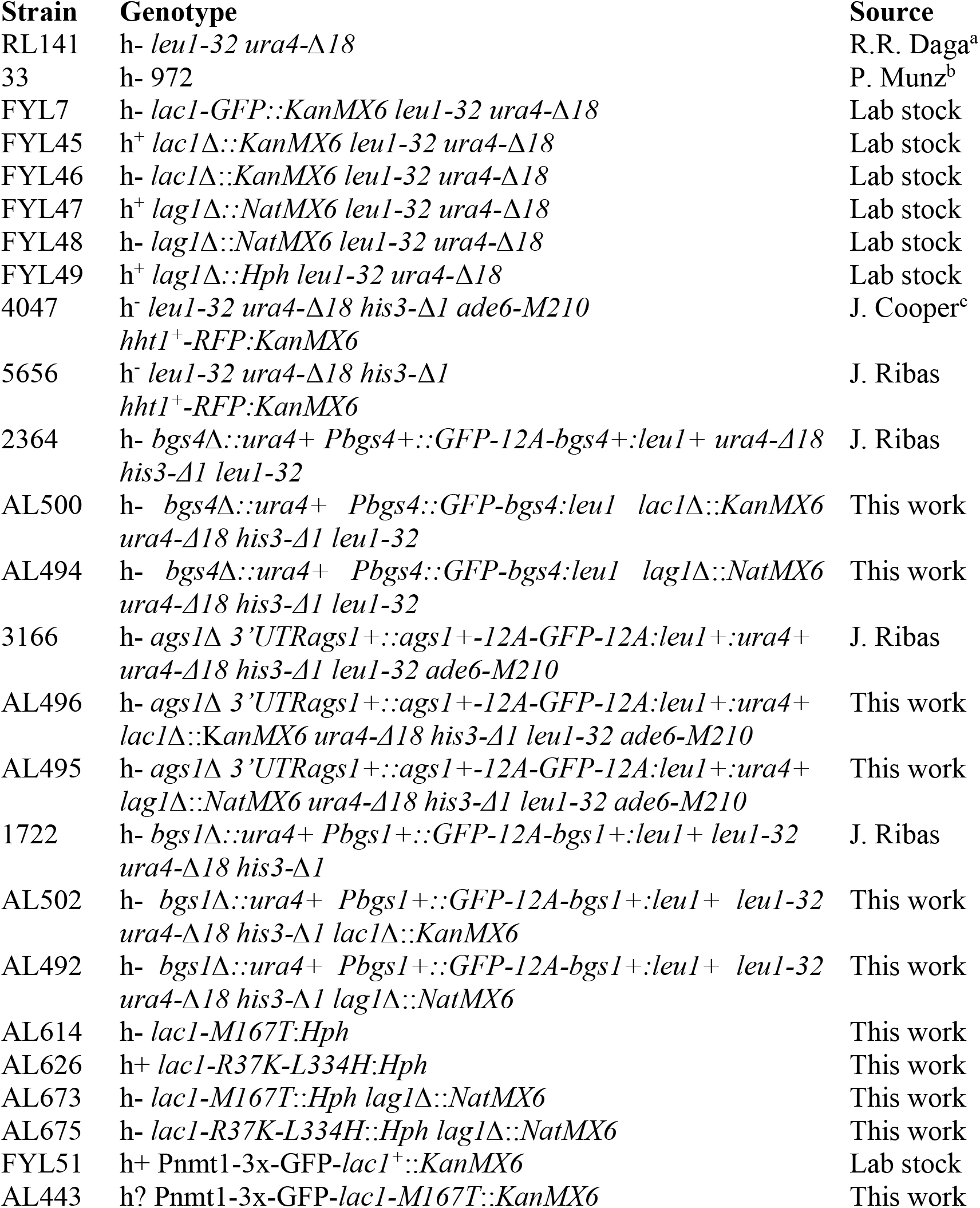
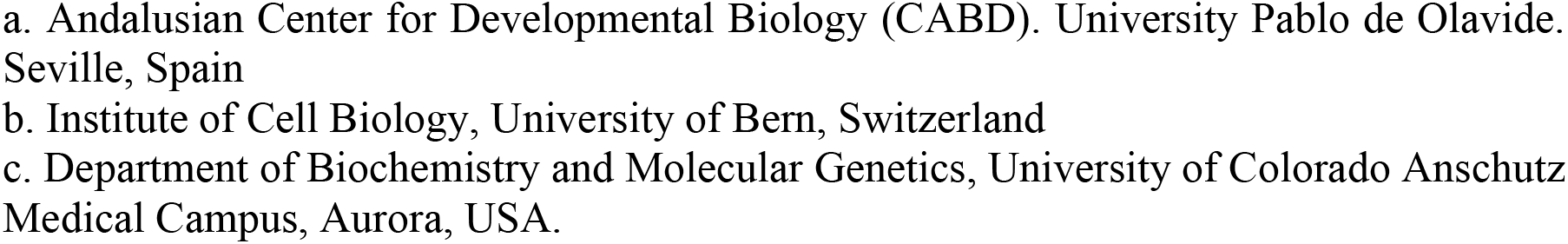
Strains used in this study.

| Strain | Genotype | Source |
|---|---|---|
| RL141 | h- *leu1-32 ura4-Δ18* | R.R. Daga<sup>a</sup> |
| 33 | h- 972 | P. Munz<sup>b</sup> |
| FYL7 | h- *lac1-GFP::KanMX6 leu1-32 ura4-Δ18* | Lab stock |
| FYL45 | h<sup>+</sup> *lac1Δ::KanMX6 leu1-32 ura4-Δ18* | Lab stock |
| FYL46 | h- *lac1Δ::KanMX6 leu1-32 ura4-Δ18* | Lab stock |
| FYL47 | h<sup>+</sup> *lag1Δ::NatMX6 leu1-32 ura4-Δ18* | Lab stock |
| FYL48 | h- *lag1Δ::NatMX6 leu1-32 ura4-Δ18* | Lab stock |
| FYL49 | h<sup>+</sup> *lag1Δ::Hph leu1-32 ura4-Δ18* | Lab stock |
| 4047 | h<sup>-</sup> *leu1-32 ura4-Δ18 his3-Δ1 ade6-M210 hht1<sup>+</sup>-RFP:KanMX6* | J. Cooper<sup>c</sup> |
| 5656 | h<sup>-</sup> *leu1-32 ura4-Δ18 his3-Δ1*<br>*hht1<sup>+</sup>-RFP:KanMX6* | J. Ribas |
| 2364 | h- *bgs4Δ::ura4+ Pbgs4+::GFP-12A-bgs4+:leu1+ ura4-Δ18 his3-Δ1 leu1-32* | J. Ribas |
| AL500 | h- *bgs4Δ::ura4+ Pbgs4::GFP-bgs4:leu1 lac1Δ::KanMX6 ura4-Δ18 his3-Δ1 leu1-32* | This work |
| AL494 | h- *bgs4Δ::ura4+ Pbgs4::GFP-bgs4:leu1 lag1Δ::NatMX6 ura4-Δ18 his3-Δ1 leu1-32* | This work |
| 3166 | h- *ags1Δ 3'UTRags1+::ags1+-12A-GFP-12A:leu1+:ura4+ ura4-Δ18 his3-Δ1 leu1-32 ade6-M210* | J. Ribas |
| AL496 | h- *ags1Δ 3'UTRags1+::ags1+-12A-GFP-12A:leu1+:ura4+ lac1Δ*::KanMX6 *ura4-Δ18 his3-Δ1 leu1-32 ade6-M210* | This work |
| AL495 | h- *ags1Δ 3'UTRags1+::ags1+-12A-GFP-12A:leu1+:ura4+ lag1Δ::NatMX6 ura4-Δ18 his3-Δ1 leu1-32 ade6-M210* | This work |
| 1722 | h- *bgs1Δ::ura4+ Pbgs1+::GFP-12A-bgs1+:leu1+ leu1-32 ura4-Δ18 his3-Δ1* | J. Ribas |
| AL502 | h- *bgs1Δ::ura4+ Pbgs1+::GFP-12A-bgs1+:leu1+ leu1-32 ura4-Δ18 his3-Δ1 lac1Δ::KanMX6* | This work |
| AL492 | h- *bgs1Δ::ura4+ Pbgs1+::GFP-12A-bgs1+:leu1+ leu1-32 ura4-Δ18 his3-Δ1 lag1Δ::NatMX6* | This work |
| AL614 | h- *lac1-M167T:Hph* | This work |
| AL626 | h+ *lac1-R37K-L334H:Hph* | This work |
| AL673 | h- *lac1-M167T::Hph lag1Δ::NatMX6* | This work |
| AL675 | h- *lac1-R37K-L334H::Hph lag1Δ::NatMX6* | This work |
| FYL51 | h+ Pnmt1-3x-GFP-*lac1<sup>+</sup>::KanMX6* | Lab stock |
| AL443 | h? Pnmt1-3x-GFP-*lac1-M167T::KanMX6* | This work |

a. Andalusian Center for Developmental Biology (CABD). University Pablo de Olavide. Seville, Spain
b. Institute of Cell Biology, University of Bern, Switzerland
c. Department of Biochemistry and Molecular Genetics, University of Colorado Anschutz Medical Campus, Aurora, USA.

For the tetrad dissection assay, cells were grown at 25°C or 30°C in SPA media. Tetrads segregants were selected on YES media supplemented with Nourseothiricin (clonNAT) at a final concentration of 100 μg/ml, G418 at a final concentration of 100 μg/ml and Hygromycin B at a final concentration of 300 μg/ml.

To assay the rate of cell proliferation on plates, cells were grown overnight in YES media at 25°C and adjusted to an A_595_ of 0.5. Ten-fold serial dilutions were spotted onto YES supplemented with Phloxine B (final concentration of 5 µg/ml) and incubated at 25°C, 20°C, 32°C, or 36°C for 3 days.

### Microscopy analysis

For the visualization of septa, cells were stained with the fluorochrome calcofluor. Briefly, 200 µL of early log-phase culture was centrifuged (1 min at 2,500 rpm). Following the removal of the supernatant, the cell pellet was resuspended in 10 µL of calcofluor white diluted to a final concentration of 50 µg/mL. Subsequently, a 1.6 µL aliquot of the cell suspension was placed on a microscope slide, and the coverslip was placed over the drop. In this case, images were taken on a LEICA DMi8 equipped with an objective lens (HCX PL APO 1003/1.40OIL PH3 CS), a Hamamatsu camera, and Application Suite X (LAS X) software. The number of septated cells was scored at each time (n > 1,000).

The analysis of cell wall and septum polymers comparing the fluorescence intensity in two strains simultaneously was performed essentially as described [16]. Equal amounts of early log-phase *hht1^+^-RFP* (control wild-type, nucleus labeling) and wild type, *lac1*Δ, or *lag1*Δ cells were mixed, washed with PBS (3,000 g for 1 min), suspended in 500 µL PBS containing 50 μg/mL calcofluor (Fluorescent Brightener 28, Sigma-Aldrich), or 1 mg/mL aniline blue (Sigma-Aldrich), and visualized for fluorescence. Other mixture of cells was suspended in 500 µL PBS containing 20 μg/mL FITC-conjugated lectin from *Bandeiraea simplicifolia* (Sigma-Aldrich, #L2895), and incubated at 28°C for 20 min in the dark with shaking. The cells were washed three times with PBS (3,000 g for 1 min), resuspended in 10 μL PBS, and visualized for FITC fluorescence.

To analyze GFP-Lac1 localization, cells were grown in early log-phase in liquid YES media at 25°C with shaking for 18-24 h and then, shifted to 36°C for 6 h. Samples were collected by centrifugation (1 min at 2,500 rpm) at each time point (**Figure 4A, B**) and placed on a microscope slide. Images were taken on a LEICA DMi8 equipped with an objective lens (HCX PL APO 1003/1.40OIL PH3 CS), L5 (GFP) filter, a Hamamatsu camera, and Application Suite X (LAS X) software. The number of cells with GFP-Lac1 mislocalized was scored at each time (n= 400).

Filipin staining is described in [54]. Briefly, 200 µL of cell cultures were incubated with 1.2 µL filipin (from a stock of 30 µg/ml) for 10 minutes. Samples were washed with 1 mL of PBS 1X and resuspend in 50 µL of PBS. Imaging was performed on a LEICA DMi8 equipped with an objective lens (HCX PL APO 1003/1.40OIL PH3 CS), a Hamamatsu camera, and Application Suite X (LAS X) software.

### Time-lapse microscopy and analysis of septal dynamics

For **Figure 1A, 4A** and **Supplementary Figure 3**, a Leica DM8000B microscope equipped with an objective lens (HCX PL APO 100×/1.40 OIL PH3 CS), a DFC350FX camera, and LAS AF software was used. Temperature was stably controlled during imaging at 25°C.

For **Figure 1C, D, and E**, 16-bits images were acquired using a fluorescence microscope (model DM RXA; Leica), a PL APO 63×/1.32 oil PH3 objective, a digital camera (modelDFC350FX; Leica), and CW4000 cytoFISHsoftware (Leica). For figure composition, the grey 16-bits images were colored and converted to 8-bits with ImageJ software (National Institutes of Health), and processed (brightness, contrast, and/or sharpness) with Adobe Photoshop CS2 software. Fluorescence intensity analysis of calcofluor, aniline blue, and FITC-conjugated lectin staining of original 16-bits images, was made with ImageJ software by placing a fixed square on the cell poles and measuring total and maximal fluorescence, and subtracting the background fluorescence value. The values from 7 independent experiments were normalized to the internal control WT-RFP value, and set to 1. The number of cells analyzed in each case is indicated in the figure legend.

For **Supplementary Figure 2**, the analysis of calcofluor and aniline blue fluorescence intensity of the septum, was made with the Line scan function of ImageJ software, and using the same original 16-bits images acquired in **Figure 1C, D, and E**. The fluorescence intensities of the corresponding septum were analyzed by drawing a line along all the septum length, and measuring every 0.1 µm the intensities of pixels of calcofluor or aniline blue stain fluorescence through the plot profile function of the Image J software. The number of cells analyzed in each case is indicated in the figure legend. Values are shown as arbitrary units (A.U.).

Fluorescence images in **Figure 2A** were acquired from early log-phase cells in the exponential growth phase using a Zeiss Axio Observer 7 inverted microscope, equipped with Alpha Plan-Apochromat 100×/1.46 Oil DIC lenses, and coupled with a Spinning Disk Confocal Yokogawa CSU-W1 head. The system was equipped with solid-state excitation lasers [50 mW, 488 nm (GFP)], and a DBP emission filter (525/50) from 3i (Intelligent Imaging Innovations). Device control and image acquisition were performed using SlideBook 6 software.

Time-lapse imaging for **Figure 2C** was performed with a spinning disk confocal microscope (IX-81; Olympus; CoolSnap HQ2 camera, Plan Apochromat 100×, 1.4 NA objective [Roper Scientific]). Images were acquired and analyzed with Metamorph software (Molecular Devices), ImageJ (National Institutes of Health), and Softworx (Applied Precision). Each time point comprised seven z-sections spaced 0.5 µm apart over a 512 × 512 pixel field, and time points were collected every 2 min (122.97 s) for 91 frames (186 min) across 48 stage positions. Images were analyzed as maximum-intensity projections of the z-stack. For imaging, early log-phase cells were placed on microslide wells (Ibidi #80821) coated with soybean lectin (Sigma #1395), and grown in YES medium supplemented as required.

To quantify septal fluorescence in **Figure 2D**, a 5 px bar was drawn across the division plane in each frame, with background taken from a zone with no cells. One trace was obtained per dividing cell for the full 91 time points.

For alignment of the onset of septal accumulation, because cells enter septation at different times within a recording, each trace was aligned on its own onset before averaging. Traces were smoothed with a five-point moving average, the frame of steepest increase was located, and the search was then walked backwards to the last local minimum preceding that increased; the onset (t = 0) was defined as the first frame at which fluorescence exceeded the minimum of 5 % of the subsequent rise amplitude. Defining the onset relative to the local minimum, rather than to the start of the recording, is necessary because some cells still carry the previous septum when imaging begins. Aligned traces were averaged from 10 frames before to 90 frames after the onset; means were plotted with the standard deviation across cells, and frames falling outside a given recording were excluded from that average.

Septal residence time in **Figure 2E** was calculated as the full time Ags1, Bgs1 and Bgs4 localized at the septum until cell separation. The residence time was considered the interval between those two frames, converted to minutes from the 2 min per frame. Residence times were compared between strains by one-way ANOVA followed by Tukey’s honest significant difference test.

### Transmission electron microscopy

Cells were grown to early log phase in YES medium at 25°C and fixed by adding an equal volume of 4% (v/v) glutaraldehyde to the culture, giving a final concentration of 2% (v/v), for 2 h at 25°C with shaking. Samples were post-fixed in 2% (v/v) glutaraldehyde with microwave assistance (700 W, 40 s, 25°C, under vacuum), washed in 0.1 M sodium cacodylate buffer pH 7.2, and stored in the same buffer until further processing.

For processing, samples were washed twice in distilled water and incubated in 1.5% (w/v) potassium permanganate for a total of 5 min, with microwave pulses (700 W, 40 s, 25°C, under vacuum) applied at the beginning and at the end of the incubation. After repeated washes in distilled water until the pellet was clear, samples were incubated in 2% (w/v) uranyl acetate for a total of 30 min with agitation, including two microwave pulses (700 W, 40 s, 25°C, under vacuum), and washed three times in distilled water. Pellets were embedded in 6% (w/v) low-melting-point agarose and trimmed into blocks of approximately 1 mm³. Blocks were dehydrated through graded acetone series (50%, 70%, 90%, 100% and 100%), each step consisting of one microwave pulse (700 W, 40 s, 25°C, without vacuum), followed by incubation to a total of 10 min per step.

Dehydrated blocks were infiltrated with Spurr’s resin through successive acetone:Spurr mixtures of 3:1, 1:1 and 1:3, each for 30 min with agitation, and including two microwave pulses (400 W, 5 min, 35°C, under vacuum). Samples were then transferred to 100% Spurr’s resin, which was applied as one microwave pulse (400 W, 5 min, 35°C, under vacuum), followed by two successive changes of 8 h and 8–10 h with agitation at 25°C and a final change with two microwave pulses (400 W, 5 min, 35°C, under vacuum). Resin was polymerized for 48 h at 60°C.

Ultrathin sections of 70 nm were cut on a Leica UC7 ultramicrotome, and collected on copper grids. Grids were post-stained with 2% (w/v) uranyl acetate for 30 min followed by lead citrate for 10 min, and sections were examined in a Zeiss Libra 120 transmission electron microscope operated at 80 kV.

### Enzyme preparation and β-(1,3)-glucan synthase assay

Membrane extracts and *in vitro* β-(1,3)-glucan synthase assays were performed essentially as described [20]. Membrane extracts were obtained from early log-phase cells grown in YES medium at 28°C. The cells were washed with buffer A (50 mM Tris-HCl, pH 7.5, 1 mM EDTA, and 1 mM β-mercaptoethanol), suspended in 150 μL of buffer A containing 300 μM GTPγS to preserve enzyme activity, and broken with glass beads in a FastPrep FP120 (MP Biomedicals, Thermo Scientific; 20 s at a speed of 6.0 and a temperature of 4 °C). Cell debris was removed by low-speed centrifugation (1500 g, 5 min, 4°C) and the supernatant was centrifuged at 48,000 g for 30 min at 4°C. Membrane extracts were resuspended in the same buffer A containing 33% glycerol and 300 μM GTPγS, and stored at −80°C. All enzymatic assays (150 μm GTPγS and 15-20 μg of protein) were carried out in duplicate, and the values were calculated from nine independent cell cultures.

Because the enzymatic specific activity depends on the obtained membrane preparation, every experiment included a wild-type control extract processed in parallel, and activities were reported as the ratio to the wild-type value of the same experiment. Nine independent experiments were analyzed. Ratios were log-transformed; each mutant was compared with its wild-type control by a one-sample t-test and the two *lac1*Δ and *lag1*Δ mutants were compared with each other by a paired t-test.

### Lac1 mutagenesis

Template DNA for PCR mutagenesis containing 400 bp of DNA upstream and 100 bp of DNA downstream of the Lac1-GFP-KanMX6 cassette was amplified from genomic DNA from strain FYL7 using primers Lac1-400_F and Lac1_+100_R, and cloned into the pGemT-easy vector (Promega #A1380). Random mutagenesis by PCR using BioTaq DNA polymerase (Bioline) in decreased dATP concentrations was carried out using primers Lac1-400_F (CCAGCAATATGCTCCTGAGGTACC) and Lac1_+100_R (CTCGTTAGTGATTGGAATTATGCG). The resulting DNA was transformed into *lag1Δ* cells (FYL49) at 25°C, grown for 24 h under non-selective conditions, and cells were replicated onto plates containing 100 mg/mL geneticin (G418) to select for transformants. Colonies were replicated on YES plates at 25°C, 32°C and 36°C, and colonies displaying impaired growth at 32°C and/or 36°C but not 25°C were isolated. Mutations were analyzed via DNA sequencing of *lac1*. Candidate alleles were amplified and integrated into strain RL141, and genetic crosses with strain FYL49 confirmed that mutant phenotypes were linked to these mutations. To remove the *GFP-KanMX6* cassette, we generated a strain with the *HphMX6* marker 123 bp downstream of *lac1* (out of the 3′UTR) through homologous recombination, using the primers Lac1-3UTR-Tag-FW (TAGAGCAGTATCCATAGTCAACGTAAATTCATATTGTGAGGTAATT) and Lac1-3UTR-TagRV (TTTGCCCCTTAGTTTTTTCCTCTAATTTTTAAAAAAACCACAACTA). Genomic DNA from this strain was amplified by PCR with the primers Lac1-3UTR-Check-FW (CGAGCACTTTTCCTTACCTTGTGAC) and RV (GCAAAGAATTTTAAACACAATGGAC) and used to make a marker switch in the strains containing the *lac1-ts-GFP-KanMX6*.

### Lipid analysis

In vivo labeling of sphingolipids with [³H]dihydrosphingosine (DHS) was carried out as described [46,56]. Briefly, cells were grown overnight in YES media supplemented with 225 mg/L adenine, histidine, lysine, uracil and lysine hydrochloride at 25°C, preincubated at 36°C for 1 h and then labeled with ^[³H]^DHS at 36°C for 2 h. Radiolabeled lipids were extracted with chloroform–methanol–water (10:10:3, v/v/v) and analyzed by TLC using solvent system I: chloroform–methanol–4.2 N ammonium hydroxide (9:7:2, v/v/v). For ceramide analysis, the fractions containing ceramides were collected by scraping the silica gel from the TLC plate and eluting the lipids with chloroform–methanol (1:1, v/v). Ceramides were then analyzed by TLC using solvent system II: chloroform–methanol–4.2 N ammonium hydroxide (40:10:1, v/v/v). Radiolabeled lipids were visualized and quantified on a Typhoon FLA-7000 system (GE Healthcare, Chicago, IL, USA). Three independent replicates were analyzed per strain.

### Structural models and substrate placement

The AlphaFold model of *Schizosaccharomyces pombe* Lac1 (UniProt O59735; model version 6) was used as the starting protein structure. The model was structurally aligned in PyMOL to the *Saccharomyces cerevisiae* Lac1 subunit of the C_26_-CoA-bound Lac1-Lip1 complex (PDB 8IZD) [57]. The experimentally observed C_26_-CoA coordinates were transferred to the *S. pombe* model through the corresponding rigid-body transformation. The membrane-oriented 8IZD structure provided by the Protein Data Bank of Transmembrane Proteins (PDBTM) was used as a reference to orient the Lac1 model with the membrane normal aligned along the z axis. Residues 58-353 of *S. pombe* Lac1 were retained for MD and capped with N-terminal acetyl and C-terminal N-methylamide groups. Each simulation system contained one Lac1 molecule and one C_26_-CoA molecule.

### Membrane system and molecular parameters

The protein–substrate complex was embedded in a pre-equilibrated yeast endoplasmic reticulum membrane obtained from Limonada (LIM4_yeast_er1; er1-200ns.pdb) [60]. Lipids with any heavy atom within 0.24 nm of a protein or substrate heavy atom were removed, leaving 229 of the original 300 lipids: 60 DYPC, 44 YOPC, 32 POPI, 24 PYPI, 15 DYPE, 14 YOPE, 13 ergosterol, 9 YOPA, 10 YOPS and 8 POPS. The two leaflets contained 116 and 113 lipids, and the initial box dimensions were 9.76473 × 9.76473 × 10.00000 nm. System preparation and MD simulations used GROMACS 2026.3 [61] and the charmm36-feb2026_cgenff-5.0 force-field distribution, ported using charmm2gmx [62] and comprising CHARMM36m protein, CHARMM36 lipid and CGenFF parameters [63–65]. C_26_-CoA parameters were derived from the C_22_-CoA model [66] by extending the acyl chain by four carbons, with a net ligand charge of −4e. Additional bonded parameters were converted from CHARMM to GROMACS functional forms and units and incorporated into the ligand topology. Water was represented by the CHARMM TIP3P model. Within 1.25 nm of the membrane midplane, waters were retained only if they were within 0.45 nm of protein or C_26_-CoA polar atoms or within 0.50 nm of H228, H229, D256 or D259. The final system contained 18,974 water molecules, 168 Na⁺ ions and 86 Cl⁻ ions; the ions provided charge neutralization and approximately 0.15 M NaCl. Neutral histidine tautomers were assigned as HSE (Nε2-protonated) at H134, H186, H189, H228, H243 and H292, and HSD (Nδ1-protonated) at H229 and mutant H334.

### WT equilibration and mutant construction

The solvated WT system underwent three stages of steepest-descent energy minimization, with maximum-force convergence thresholds of 1000, 250 and 100 kJ mol⁻¹ nm⁻¹, respectively. Protein backbone and C_26_-CoA heavy atoms were initially restrained with force constants of 1000 and 500 kJ mol⁻¹ nm⁻², respectively. The second stage retained only backbone restraints at 500 kJ mol⁻¹ nm⁻², and the third was unrestrained. Three WT replicates were equilibrated at 298.15 K for 200 ps under NVT conditions and 2.5 ns under NPT conditions, with restraints of 500 kJ mol⁻¹ nm⁻² on protein backbone and C_26_-CoA heavy atoms. Both restraints were reduced to 100 kJ mol⁻¹ nm⁻² for a further 250 ps of NPT equilibration, followed by 500 ps of unrestrained NPT relaxation. M167T and L334H models were generated in PyMOL from each equilibrated WT replicate, retaining its substrate, membrane, solvent, ions and simulation box. Mutant protein topologies were regenerated. Each mutant was energy-minimized and equilibrated at 298.15 K for 100 ps under NVT conditions, with restraints of 100 kJ mol⁻¹ nm⁻² in each Cartesian direction on protein atoms named N, CA, C and O and C_26_-CoA heavy atoms, referenced to the minimized coordinates. This was followed by 250 ps of unrestrained NPT equilibration. All WT and mutant replicates subsequently underwent 100 ps of NVT and 400 ps of NPT equilibration at 309.15 K (36°C). Initial velocities were generated separately for the initial WT and mutant NVT stages from a Maxwell distribution at 298.15 K, using a different random seed for each replicate. The remaining equilibration stages inherited coordinates, velocities and box dimensions from the preceding checkpoint without regenerating velocities. Integration, bond-constraint, nonbonded-interaction and thermostat settings were as described below for production simulations; NPT stages also used the same barostat settings.

### Production simulations

Three independent 200 ns production simulations were performed for each genotype at 309.15 K and 1 bar. Simulations continued from the final equilibration checkpoints without regenerating velocities or applying positional restraints. A 2 fs leapfrog timestep and three-dimensional periodic boundary conditions were used. Solute and solvent were coupled separately using the velocity-rescaling thermostat [67] with a coupling time of 1 ps. Pressure was controlled by semi-isotropic stochastic cell rescaling (C-rescale) [68], with a coupling time of 5 ps and a compressibility of 4.5 × 10⁻⁵ bar⁻¹ in both coupled directions. Bonds involving hydrogen in non-water molecules were constrained using LINCS [69], with an expansion order of 4 and two iterations; water geometry was constrained using SETTLE [70]. Verlet neighbor lists were used, with automatic adjustment of the update interval and buffer. Electrostatic interactions were calculated using particle-mesh Ewald [71] with a real-space cutoff of 1.2 nm, fourth-order interpolation and a Fourier grid spacing of 0.12 nm. Lennard–Jones forces were switched to zero between 1.0 and 1.2 nm, without long-range dispersion correction. Compressed coordinates, energies and log entries were saved every 10 ps.

### Trajectory analysis

Trajectories were analyzed at 100 ps intervals. Protein molecules were made whole under periodic boundary conditions before fitting. Catalytic-core backbone RMSD was calculated for the N, CA and C atoms of H228, H229, D256 and D259 after equally weighted least-squares fitting of these atoms to the first production frame of each replicate. The distance between H228 Nε2 and the C_26_-CoA thioester carbonyl carbon (C1) was calculated using the instantaneous periodic box. Proximity occupancy was defined as the fraction of frames with a distance ≤0.45 nm. TM8 engagement was assessed for W344, L347, I348 and I351. A residue was engaged when at least one heavy-atom pair with the 48-heavy-atom CoA headgroup was separated by <0.45 nm. High engagement was defined as simultaneous engagement of at least three of the four residues. Because mutant systems were built from equilibrated WT replicates and re-equilibrated for a shorter period, TM8 engagement was summarized over the final 50 ns (150–200 ns) of each trajectory; high-engagement occupancies were calculated as time-weighted fractions over this window, assigning 0.05 ns to endpoint frames and 0.10 ns to interior frames. Residue 334–acyl-chain contacts were defined by a minimum distance <0.45 nm between all residue 334 heavy atoms, including backbone atoms, and acyl carbons at positions 13–16, numbered from the thioester carbonyl carbon as C1. Contact occupancy was calculated as the fraction of frames containing at least one contact. Catalytic-core summaries and residue 334 contact occupancies used the interval 0 ≤ t < 200 ns. Time courses were calculated separately for each trajectory using nonoverlapping, left-closed and right-open 1 ns bins. Instantaneous engaged-residue counts or binary occupancy indicators were averaged within each bin.

### Statistical analysis

A minimum of two independent biological replicates was performed for each experimental condition. For the statistical analyses, a one-way ANOVA test was performed using Prism 10 (GraphPad), followed by a Tukey post-hoc test, except for the septal fluorescence and septal residence analyses, which were performed in Python 3.11 with SciPy 1.17. For the lipid analysis and the molecular dynamics simulations, statistical analyses were performed using R 4.3.3 and its stats package (R Core Team, 2024). Each TLC replicate or MD trajectory contributed one summary value per outcome and analysis window (n = 3 per strain or genotype). Differences were assessed separately for each outcome and window using one-way ANOVA followed by Tukey’s honestly significant difference test, and Tukey-adjusted P values are reported. Cell death and septation indices were analyzed by two-way ANOVA with strain and temperature as factors, followed by Tukey’s test for comparisons against wild-type within a temperature and Šídák’s test for comparisons between temperatures within a strain; p values are reported adjusted for multiple comparisons. For the glucan synthase assay, specific activities were normalized to the specific activity of the wild-type membrane extract of the same experiment, log-transformed, and each mutant compared with its control by a one-sample t-test. Lac1 mislocalization was analyzed by ordinary two-way ANOVA with genotype and time as factors, followed by Bonferroni’s multiple comparisons test between genotypes within each time point.

## AI statement

ChatGPT (OpenAI, USA) and Claude were used to assist in writing Python scripts for running GROMACS simulations and statistical analyses in R, identifying and resolving errors, and proofreading the Methods section. The authors take full responsibility for the final code, analyses, and manuscript content.

## Author contributions

Conceptualization: I.F-P, R.L.; Methodology: K.M,; Validation: All authors; Formal analysis: K.M, K.H, K.F, R.L.; Investigation: A.S, L.G-D, K.M, K.H., J.C.G-C, M.A-G, I.F-P, R.L; Resources: K.F, J.C.G-C, J.C.R, M.A-G, I.F-P, R.L; Writing - original draft: R.L.; Writing - review & editing: A.S, K.H, J.C.G-C, J.C.R, M.A-G, I.F-P, R.L; Visualization: A.S, L.G-D, K.H, J.C.R, M.A-G, I.F-P, R.L Supervision: K.H, K.F, J.C.R, M.A-G, I.F-P, R.L; Project administration: I.F-P, R.L.; Funding acquisition: J.C.G-C, J.C.R, M.A-G., I.F-P, R.L.

## Acknowledgments

We thank members of the laboratory for advice and support. We thank R.R. Daga, P. Munz and J. Cooper for strains. We thank Derek McCusker (University of Bordeaux, France) for western blotting materials. We thank Katherina Garcia (CABD microscopy facility), and Manuel Barragan and Juan Luis Ribas (CITIUS electron microscopy service) for technical help. Laura Gómez-Delgado acknowledges support by a fellowship from Ministerio de Ciencia, Innovación y Universidades (MICIU, Spain).

## REFERENCES

1 Jorgensen P & Tyers M (2004) How cells coordinate growth and division. Current Biology 14, 1014–1027.

2 Gow NAR & Lenardon MD (2023) Architecture of the dynamic fungal cell wall. Nat Rev Microbiol 21, 248–259.

3 Davì V & Minc N (2015) Mechanics and morphogenesis of fission yeast cells. Curr Opin Microbiol 28, 36–45.

4 Jasani A, Huynh T & Kellogg DR (2020) Growth-Dependent Activation of Protein Kinases Suggests a Mechanism for Measuring Cell Growth. Genetics 215, 729–746.

5 McCusker D & Kellogg DR (2012) Plasma membrane growth during the cell cycle: unsolved mysteries and recent progress. Curr Opin Cell Biol 24, 845–851.

6 Emmerstorfer-Augustin A & Thorner J (2023) Regulation of TORC2 Function and Localization in Yeast. Annu Rev Cell Dev Biol 39.

7 Pan KZ, Saunders TE, Flor-Parra I, Howard M & Chang F (2014) Cortical regulation of cell size by a sizer cdr2p. Elife 2014, 1–24.

8 Wood E & Nurse P (2014) Sizing Up to Divide: Mitotic Cell Size Control in Fission Yeast. The Annual Review of Cell and Developmental Biology.

9 Mitchison JM & Nurse P (1985) Growth in cell length in the fission yeast Schizosaccharomyces pombe. J Cell Sci 75, 357–76.

10 Ramos M, Cortés JCG, Sato M, Rincón SA, Belén Moreno M, Clemente-Ramos JÁ, Osumi M, Pérez P & Ribas JC (2019) Two S. Pombe septation phases differ in ingression rate, septum structure, and response to F-actin loss. Journal of Cell Biology 218, 4171–4194.

11 G. Cortés JC, Ramos M, Konomi M, Barragán I, Moreno MB, Alcaide-Gavilán M, Moreno S, Osumi M, Pérez P & Ribas JC (2018) Specific detection of fission yeast primary septum reveals septum and cleavage furrow ingression during early anaphase independent of mitosis completion. PLoS Genet 14, e1007388.

12 Willet AH, McDonald NA & Gould KL (2015) Regulation of contractile ring formation and septation in Schizosaccharomyces pombe. Curr Opin Microbiol 28, 46–52.

13 García Cortés JC, Ramos M, Osumi M, Pérez P & Ribas JC (2016) The Cell Biology of Fission Yeast Septation. Microbiology and Molecular Biology Reviews 80, 779–791.

14 Ramos M, Cortés JCG, Sato M, Rincón SA, Belén Moreno M, Clemente-Ramos JÁ, Osumi M, Pérez P & Ribas JC (2019) Two S. Pombe septation phases differ in ingression rate, septum structure, and response to F-actin loss. Journal of Cell Biology 218, 4171–4194.

15 Muñoz J, Cortés JCG, Sipiczki M, Ramos M, Clemente-Ramos JA, Moreno MB, Martins IM, Pérez P & Ribas JC (2013) Extracellular cell wall β(1,3)glucan is required to couple septation to actomyosin ring contraction. Journal of Cell Biology 203, 265–282.

16 Cortés JCG, Sato M, Muñoz J, Belén Moreno M, Clemente-Ramos JA, Ramos M, Okada H, Osumi M, Durán A & Ribas JC (2012) Fission yeast Ags1 confers the essential septum strength needed for safe gradual cell abscission. Journal of Cell Biology 198, 637–656.

17 Pérez P, Cortés JCG, Cansado J & Ribas JC (2018) Fission yeast cell wall biosynthesis and cell integrity signalling. Cell Surface 4, 1–9.

18 Hochstenbach F, Klis FM, van den Ende H, van Donselaar E, Peters PJ & Klausner RD (1998) Identification of a putative alpha-glucan synthase essential for cell wall construction and morphogenesis in fission yeast. Proceedings of the National Academy of Sciences 95, 9161– 9166.

19 Katayama S, Hirata D, Arellano M, Pérez P & Toda T (1999) Fission Yeast α-Glucan Synthase Mok1 Requires the Actin Cytoskeleton to Localize the Sites of Growth and Plays an Essential Role in Cell Morphogenesis Downstream of Protein Kinase C Function. J Cell Biol 144, 1173–1186.

20 Cortés JCG, Ishiguro J, Durán A & Ribas JC (2002) Localization of the (1,3)β-D-glucan synthase catalytic subunit homologue Bgs1p/Cps1p from fission yeast suggests that it is involved in septation, polarized growth, mating, spore wall formation and spore germination. J Cell Sci 115, 4081–4096.

21 Cortés JCG, Konomi M, Martins IM, Muñoz J, Moreno MB, Osumi M, Durán A & Ribas JC (2007) The (1,3)β-D-glucan synthase subunit Bgs1p is responsible for the fission yeast primary septum formation. Mol Microbiol 65, 201–217.

22 Martín V, García B, Carnero E, Durán A & Sánchez Y (2003) Bgs3p, a putative 1,3-β-glucan synthase subunit, is required for cell wall assembly in Schizosaccharomyces pombe. Eukaryot Cell 2, 159–169.

23 Ohashi T, Fujiyama K & Takegawa K (2012) Identification of novel α1,3-galactosyltransferase and elimination of α-galactose-containing glycans by disruption of multiple α-galactosyltransferase genes in Schizosaccharomyces pombe. Journal of Biological Chemistry 287, 38866–38875.

24 Free SJ (2013) Fungal Cell Wall Organization and Biosynthesis. In Advances in Genetics pp. 33–82. Academic Press Inc.

25 García Cortés JC, Ramos M, Osumi M, Pérez P & Ribas JC (2016) The Cell Biology of Fission Yeast Septation. Microbiology and Molecular Biology Reviews 80, 779–791.

26 Curto MÁ, Butassi E, Ribas JC, Svetaz LA & Cortés JCG (2021) Natural products targeting the synthesis of β(1,3)-D-glucan and chitin of the fungal cell wall. Existing drugs and recent findings. Phytomedicine 88, 153556.

27 Cortés JCG, Curto MÁ, Carvalho VSD, Pérez P & Ribas JC (2019) The fungal cell wall as a target for the development of new antifungal therapies. Biotechnol Adv 37.

28 Downes CP & Currie RA (1998) Lipid signalling. Current Biology 8, R865–R867.

29 Lingwood D & Simons K (2010) Lipid Rafts As a Membrane-Organizing Principle. Science (1979) 327, 46–50.

30 Simons K & Ikonen E (1997) Functional rafts in cell membranes. Nature 387, 569–572.

31 Alonso A & Goñi FM (2018) The Physical Properties of Ceramides in Membranes. Annu Rev Biophys 47, 633–654.

32 Gómez-Muñoz A (1998) Modulation of cell signalling by ceramides. Biochimica et Biophysica Acta - Lipids and Lipid Metabolism 1391, 92–109.

33 Stancevic B & Kolesnick R (2010) Ceramide-rich platforms in transmembrane signaling. FEBS Lett 584, 1728–1740.

34 Lucena R, Alcaide-Gavilán M, Schubert K, He M, Domnauer MG, Marquer C, Klose C, Surma MA & Kellogg DR (2018) Cell Size and Growth Rate Are Modulated by TORC2-Dependent Signals. Current Biology 28, 196–210.e4.

35 Goñi FM & Alonso A (2006) Biophysics of sphingolipids I. Membrane properties of sphingosine, ceramides and other simple sphingolipids. Biochim Biophys Acta Biomembr 1758, 1902–1921.

36 Morad SA & Cabot MC (2013) Ceramide-orchestrated signalling in cancer cells. Nat Rev Cancer 13, 51–65.

37 Rodriguez-Gallardo S, Kurokawa K, Sabido-Bozo S, Cortes-Gomez A, Ikeda A, Zoni V, Aguilera-Romero A, Perez-Linero AM, Lopez S, Waga M, Araki M, Nakano M, Riezman H, Funato K, Vanni S, Nakano A & Muñiz M (2020) Ceramide chain length-dependent protein sorting into selective endoplasmic reticulum exit sites. Sci Adv 6.

38 Alizadeh J, da Silva Rosa SC, Weng X, Jacobs J, Lorzadeh S, Ravandi A, Vitorino R, Pecic S, Zivkovic A, Stark H, Shojaei S & Ghavami S (2023) Ceramides and ceramide synthases in cancer: Focus on apoptosis and autophagy. Eur J Cell Biol 102, 151337.

39 Chaurasia B & Summers SA (2021) Ceramides in Metabolism: Key Lipotoxic Players. Annu Rev Physiol 83, 303–330.

40 Green CD, Maceyka M, Cowart LA & Spiegel S (2021) Sphingolipids in metabolic disease: The good, the bad, and the unknown. Cell Metab 33, 1293–1306.

41 Kumar M, Singh A, Kumari S, Kumar P, Wasi M, Mondal AK, Rudramurthy SM, Chakrabarti A, Gaur NA, Gow NAR & Prasad R (2021) Sphingolipidomics of drug resistant Candida auris clinical isolates reveal distinct sphingolipid species signatures. Biochim Biophys Acta Mol Cell Biol Lipids 1866, 158815.

42 Urbanek AK, Muraszko J, Derkacz D, Łukaszewicz M, Bernat P & Krasowska A (2022) The Role of Ergosterol and Sphingolipids in the Localization and Activity of Candida albicans’ Multidrug Transporter Cdr1p and Plasma Membrane ATPase Pma1p. Int J Mol Sci 23.

43 Beeler T, Bacikova D, Gable K, Hopkins L, Johnson C, Slife H & Dunn T (1998) The Saccharomyces cerevisiae TSC10/YBR265W gene encoding 3-ketosphinganine reductase is identified in a screen for temperature-sensitive suppressors of the Ca2+-sensitive csg2Δ mutant. Journal of Biological Chemistry 273, 30688–30694.

44 Haak D, Gable K, Beeler T & Dunn T (1997) Hydroxylation of Saccharomyces cerevisiae ceramides requires Sur2p and Scs7p. Journal of Biological Chemistry 272, 29704–29710.

45 Dickson RC (2008) Thematic Review Series: Sphingolipids. New insights into sphingolipid metabolism and function in budding yeast. J Lipid Res 49, 909–921.

46 Flor-Parra I, Sabido-Bozo S, Ikeda A, Hanaoka K, Aguilera-Romero A, Funato K, Muñiz M & Lucena R (2022) The ceramide synthase subunit lac1 regulates cell growth and size in fission yeast. Int J Mol Sci 23, 1–14.

47 Nakase M, Tani M, Morita T, Kitamoto HK, Kashiwazaki J, Nakamura T, Hosomi A, Tanaka N & Takegawa K (2010) Mannosylinositol phosphorylceramide is a major sphingolipid component and is required for proper localization of plasma-membrane proteins in Schizosaccharomyces pombe. J Cell Sci 123, 1578–1587.

48 Quesada-Márquez JI, Serrano A, Alcaide-Gavilán M & Lucena R (2026) Uncoupling of nutrient sensing and cell size control by specific defects in ceramide structure. Biol Open 15.

49 Dawson K, Toone WM, Jones N & Wilkinson CRM (2008) Loss of regulators of vacuolar ATPase function and ceramide synthesis results in multidrug sensitivity in Schizosaccharomyces pombe. Eukaryot Cell 7, 926–937.

50 Ishiguro J, Saitou A, Durán A & Ribas JC (1997) cps1+, a Schizosaccharomyces pombe gene homolog of Saccharomyces cerevisiae FKS genes whose mutation confers hypersensitivity to cyclosporin A and papulacandin B. J Bacteriol 179, 7653–7662.

51 Cortés JCG, Carnero E, Ishiguro J, Sánchez Y, Durán A & Ribas JC (2005) The novel fission yeast (1,3)β-D-glucan synthase catalytic subunit Bgs4p is essential during both cytokinesis and polarized growth. J Cell Sci 118, 157–174.

52 Arasada R & Pollard TD (2014) Contractile Ring Stability in S.pombe Depends on F-BAR Protein Cdc15p and Bgs1p Transport from the Golgi Complex. Cell Rep 8, 1533–1544.

53 Arasada R & Pollard TD (2015) A role for F-BAR protein Rga7p during cytokinesis in S. pombe. J Cell Sci 128, 2259–2268.

54 Takeda T, Kawate T & Chang F (2004) Organization of a sterol-rich membrane domain by cdc15p during cytokinesis in fission yeast. Nat Cell Biol 6, 1142–1144.

55 Ramos M, Martín-García R, Curto MÁ, Gómez-Delgado L, Moreno MB, Sato M, Portales E, Osumi M, Rincón SA, Pérez P, Ribas JC & Cortés JCG (2024) Fission yeast Bgs1 glucan synthase participates in the control of growth polarity and membrane traffic. iScience 27.

56 Ikeda A, Hanaoka K & Funato K (2021) Protocol for measuring sphingolipid metabolism in budding yeast. STAR Protoc 2, 100412.

57 Xie T, Fang Q, Zhang Z, Wang Y, Dong F & Gong X (2023) Structure and mechanism of a eukaryotic ceramide synthase complex. EMBO J.

58 Nakao K, Carvalho VSD, Suganaga A, Osumi M, Tokukura M, Kakeya H, Matsuyama A, Yashiroda Y, Matsunaga S, Cortés JCG, Yoshida M, Ribas JC & Nishimura S (2026) Ectopic overproduction of cell wall glucan through membrane perturbation by an antifungal peptide theonellamide A in fission yeast. iScience 29.

59 Moreno S, Klar A & Nurse P (1991) Guide to Yeast Genetics and Molecular Biology. Methods Enzymol 194, 795–823.

60 Monje-Galvan V & Klauda JB (2015) Modeling Yeast Organelle Membranes and How Lipid Diversity Influences Bilayer Properties. Biochemistry 54, 6852–6861.

61 Abraham MJ, Murtola T, Schulz R, Páll S, Smith JC, Hess B & Lindah E (2015) Gromacs: High performance molecular simulations through multi-level parallelism from laptops to supercomputers. SoftwareX 1–2, 19–25.

62 Wacha AF & Lemkul JA (2023) charmm2gmx: An Automated Method to Port the CHARMM Additive Force Field to GROMACS. J Chem Inf Model 63, 4246–4252.

63 Klauda JB, Venable RM, Freites JA, O’Connor JW, Tobias DJ, Mondragon-Ramirez C, Vorobyov I, MacKerell AD & Pastor RW (2010) Update of the CHARMM All-Atom Additive Force Field for Lipids: Validation on Six Lipid Types. Journal of Physical Chemistry B 114, 7830–7843.

64 Vanommeslaeghe K, Hatcher E, Acharya C, Kundu S, Zhong S, Shim J, Darian E, Guvench O, Lopes P, Vorobyov I & Mackerell AD (2010) CHARMM general force field: A force field for drug-like molecules compatible with the CHARMM all-atom additive biological force fields. J Comput Chem 31, 671–690.

65 Huang J, Rauscher S, Nawrocki G, Ran T, Feig M, De Groot BL, Grubmüller H & MacKerell AD (2016) CHARMM36m: An improved force field for folded and intrinsically disordered proteins. Nat Methods 14, 71–73.

66 Stix R, Song J, Banerjee A & Faraldo-Gómez JD (2020) DHHC20 Palmitoyl-Transferase Reshapes the Membrane to Foster Catalysis. Biophys J 118, 980–988.

67 Bussi G, Donadio D & Parrinello M (2007) Canonical sampling through velocity rescaling. Journal of Chemical Physics 126.

68 Bernetti M & Bussi G (2020) Pressure control using stochastic cell rescaling. Journal of Chemical Physics 153.

69 Hess B, Bekker H, Berendsen HJC & Fraaije JGEM (1997) LINCS: A Linear Constraint Solver for Molecular Simulations John Wiley & Sons, Inc.

70 Miyamoto S & Kollman PA (1992) Settle: An analytical version of the SHAKE and RATTLE algorithm for rigid water models. J Comput Chem 13, 952–962.

71 Essmann U, Perera L, Berkowitz ML, Darden T, Lee H & Pedersen LG (1995) A smooth particle mesh Ewald method. J Chem Phys 103, 8577–8593.

