## Supplementary figures and images for "Lac1-dependent ceramide synthesis couples septum assembly to cell separation in *Schizosaccharomyces pombe*"

### Supplemental Figure 1

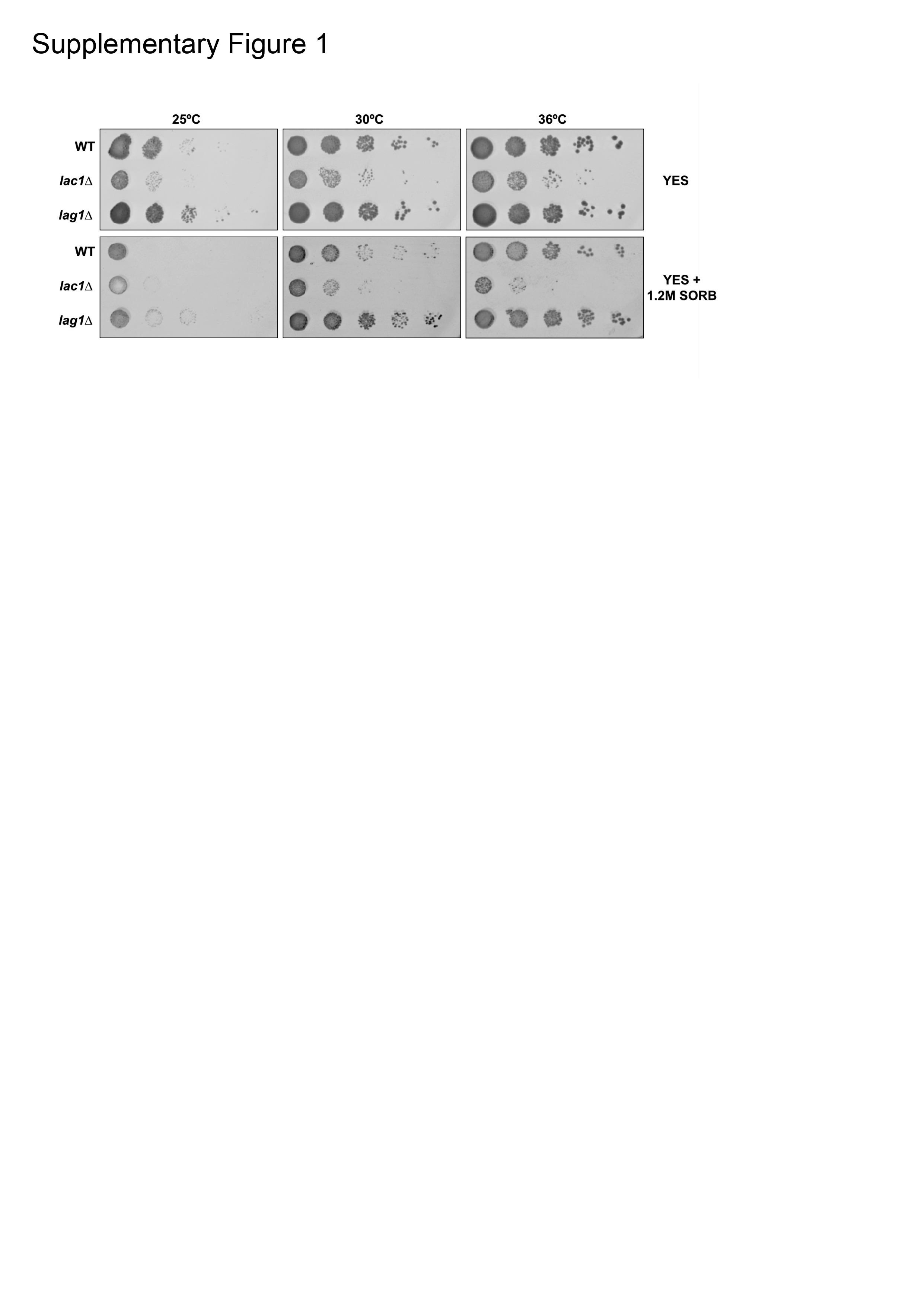

### Supplemental Figure 2

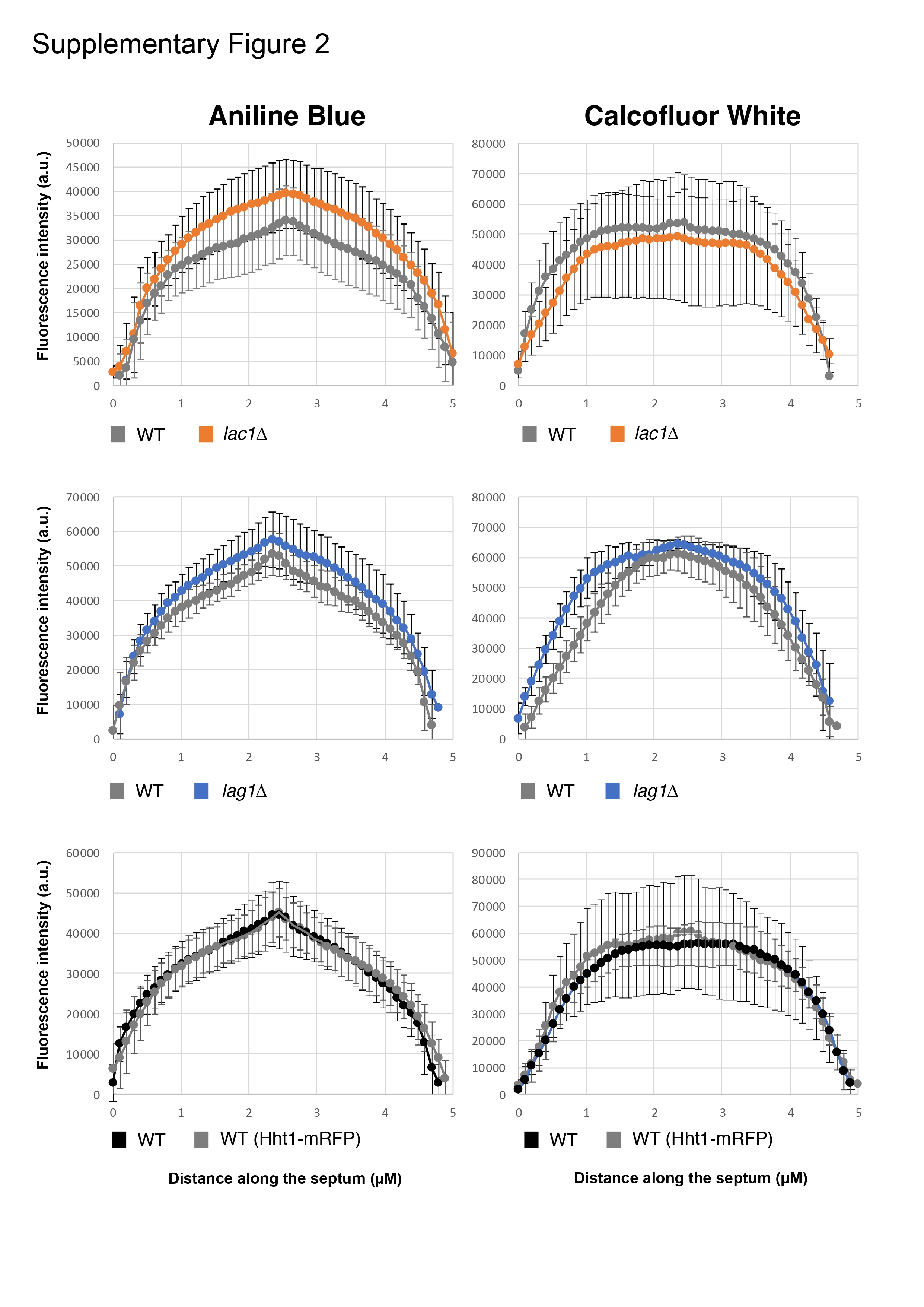

### Supplemental Figure 4

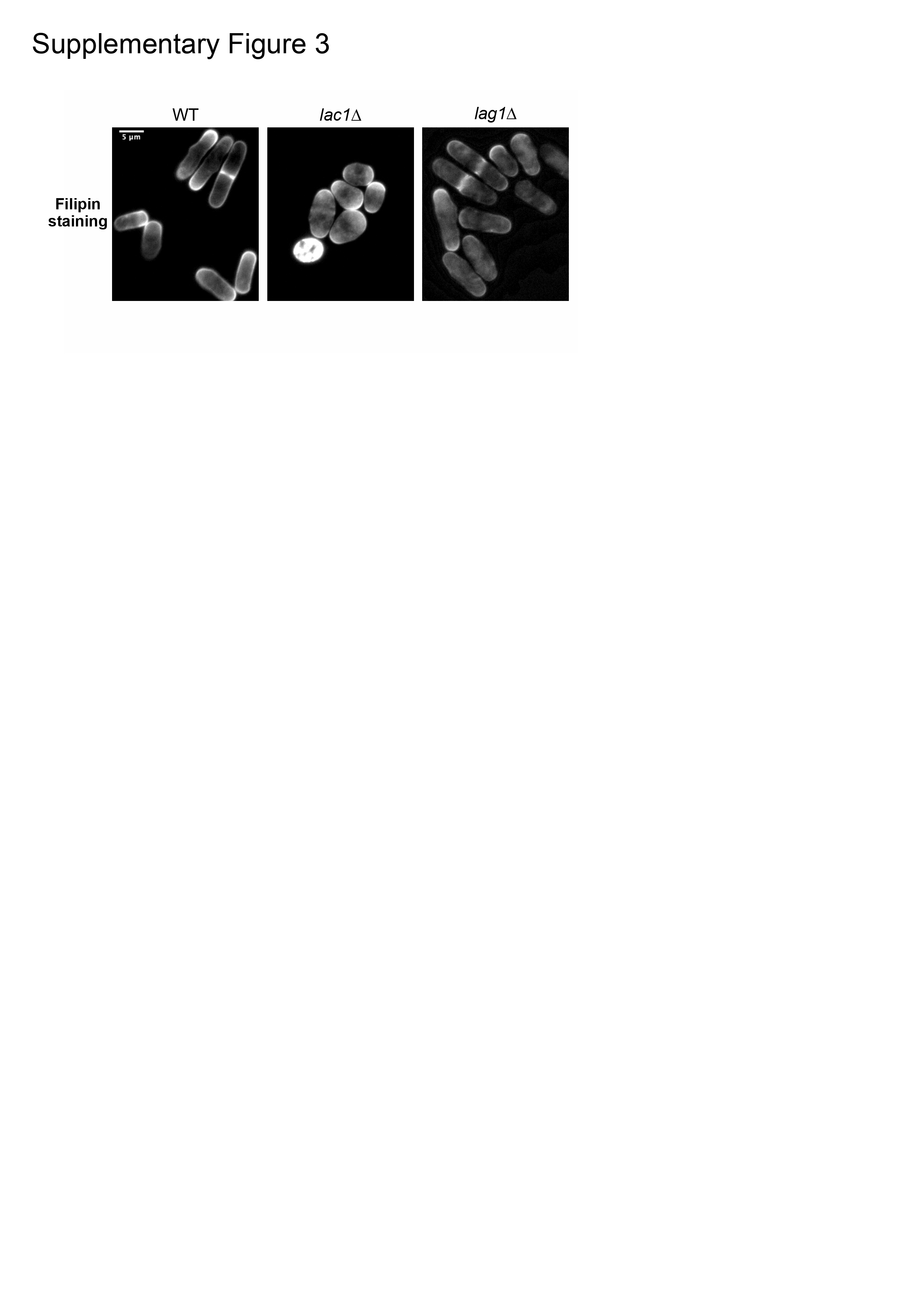

### Supplemental Figure 4

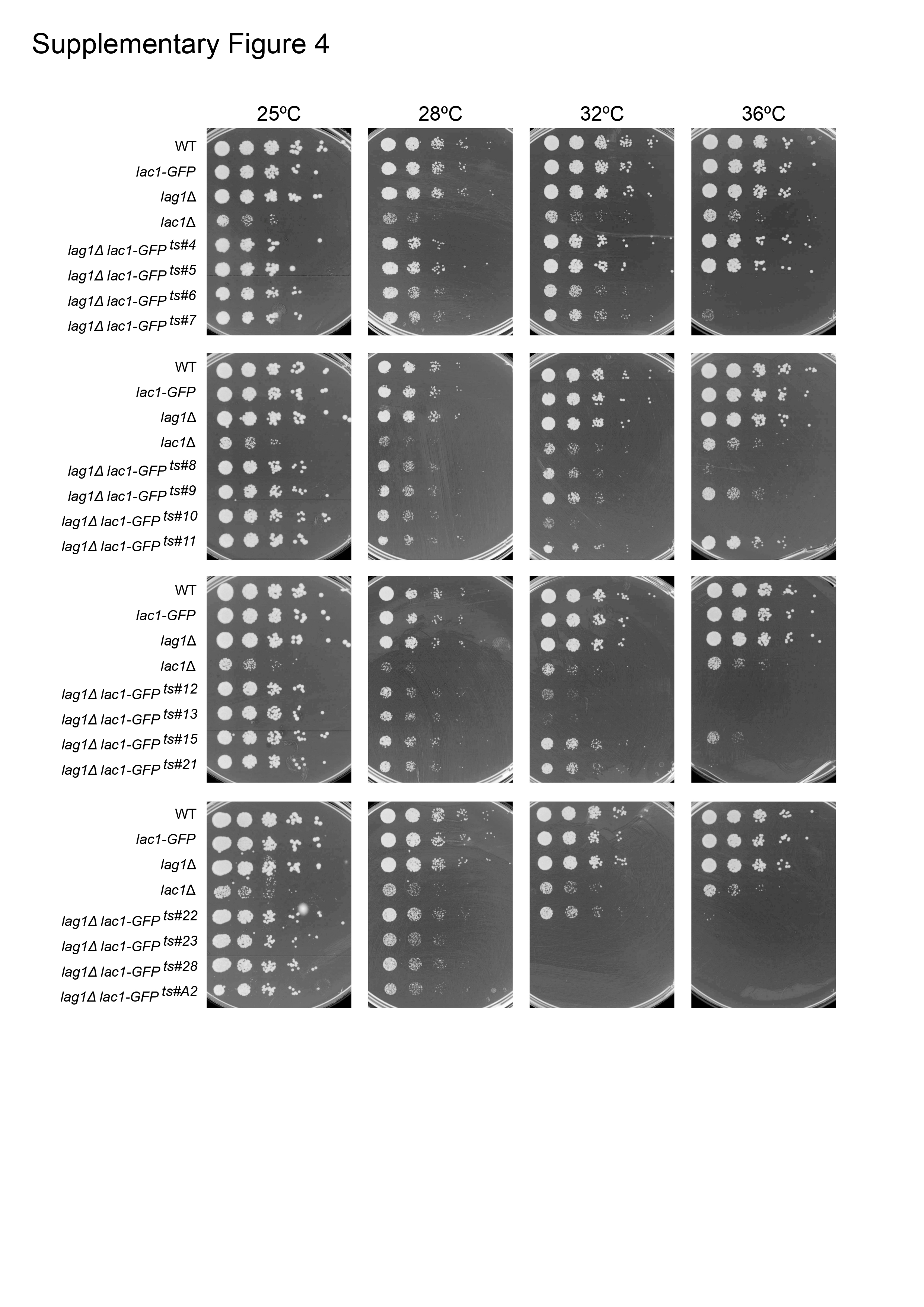

### Supplemental Figure 5

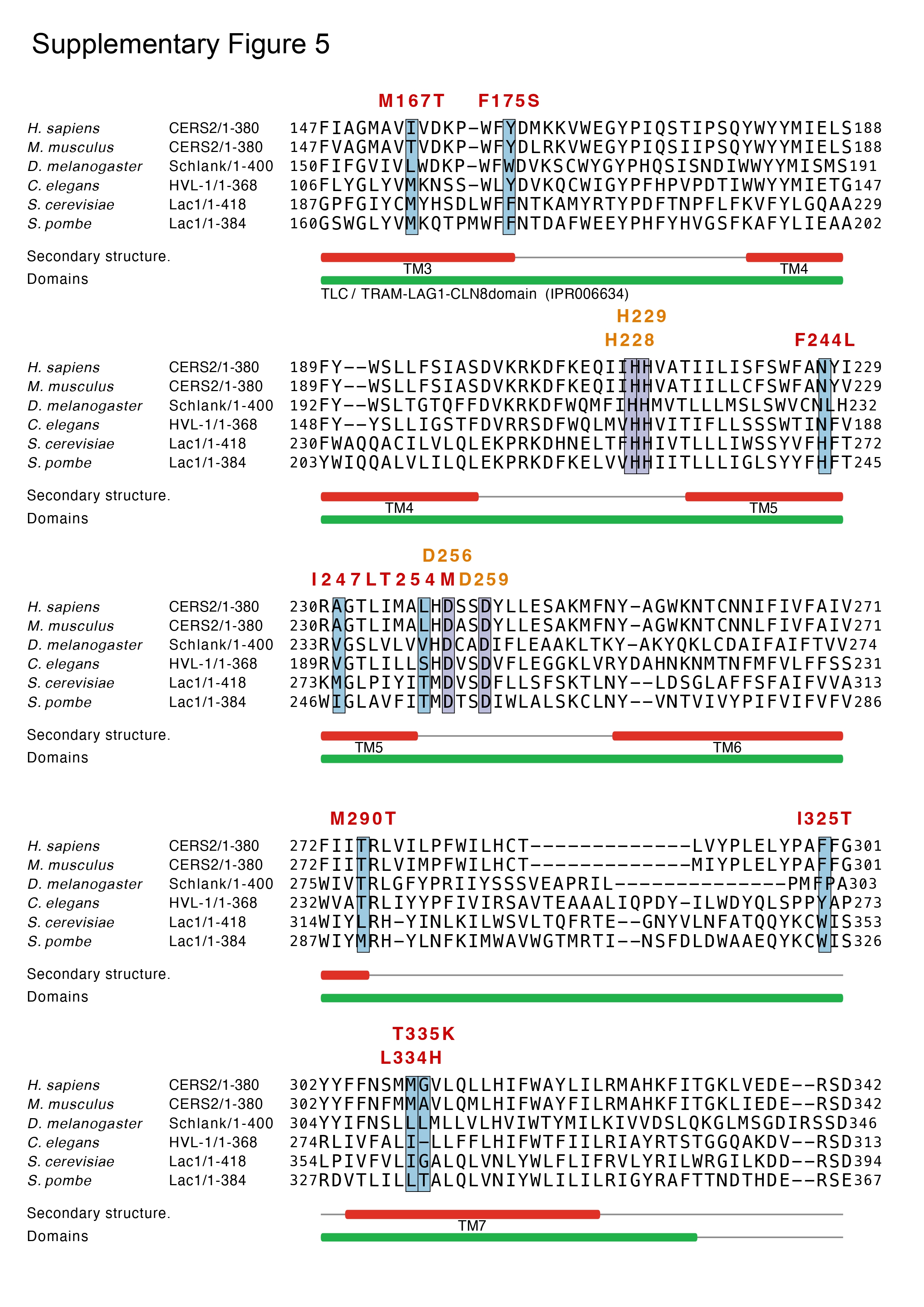

### Supplemental Figure 6

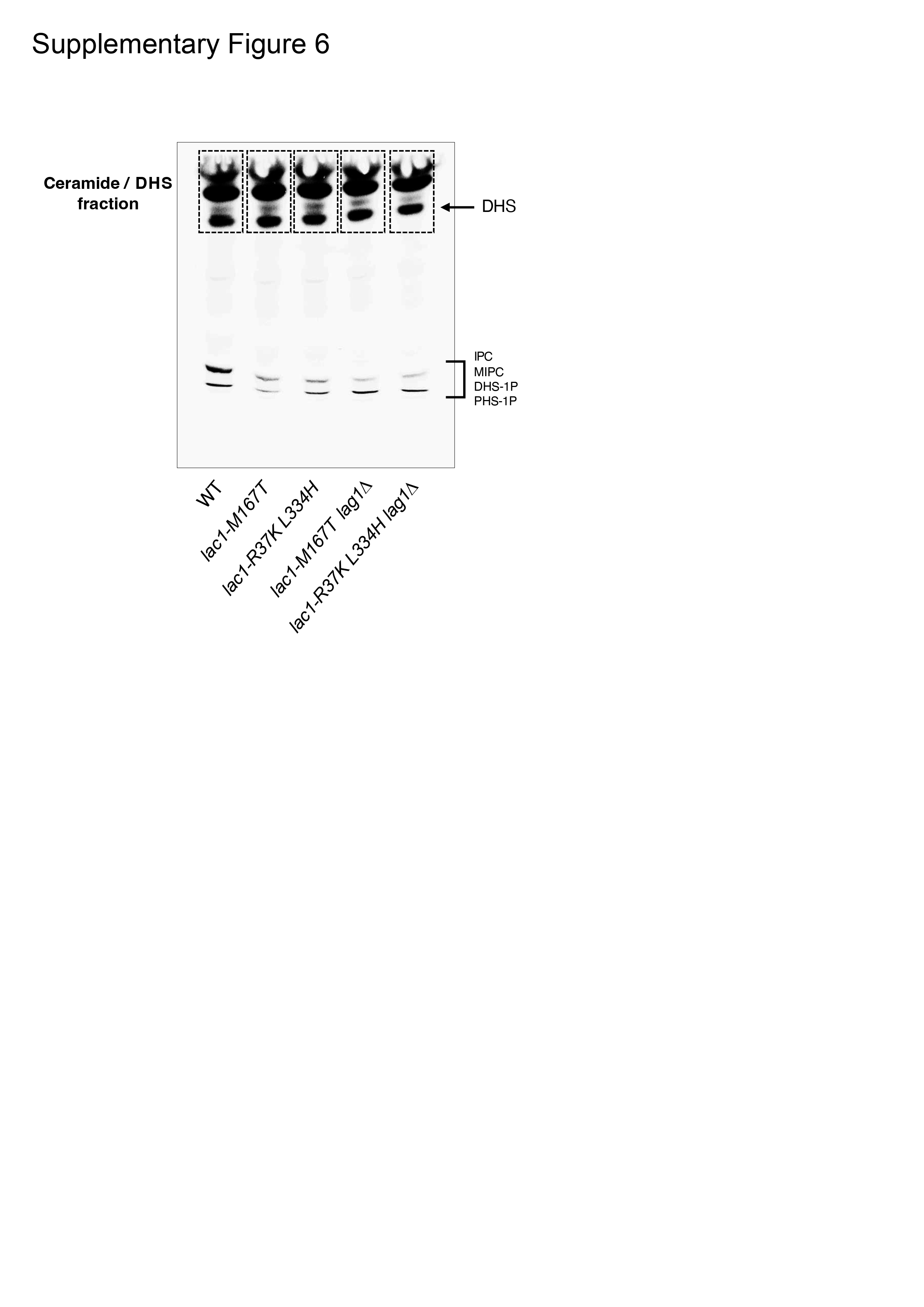

### Supplemental Figure 7

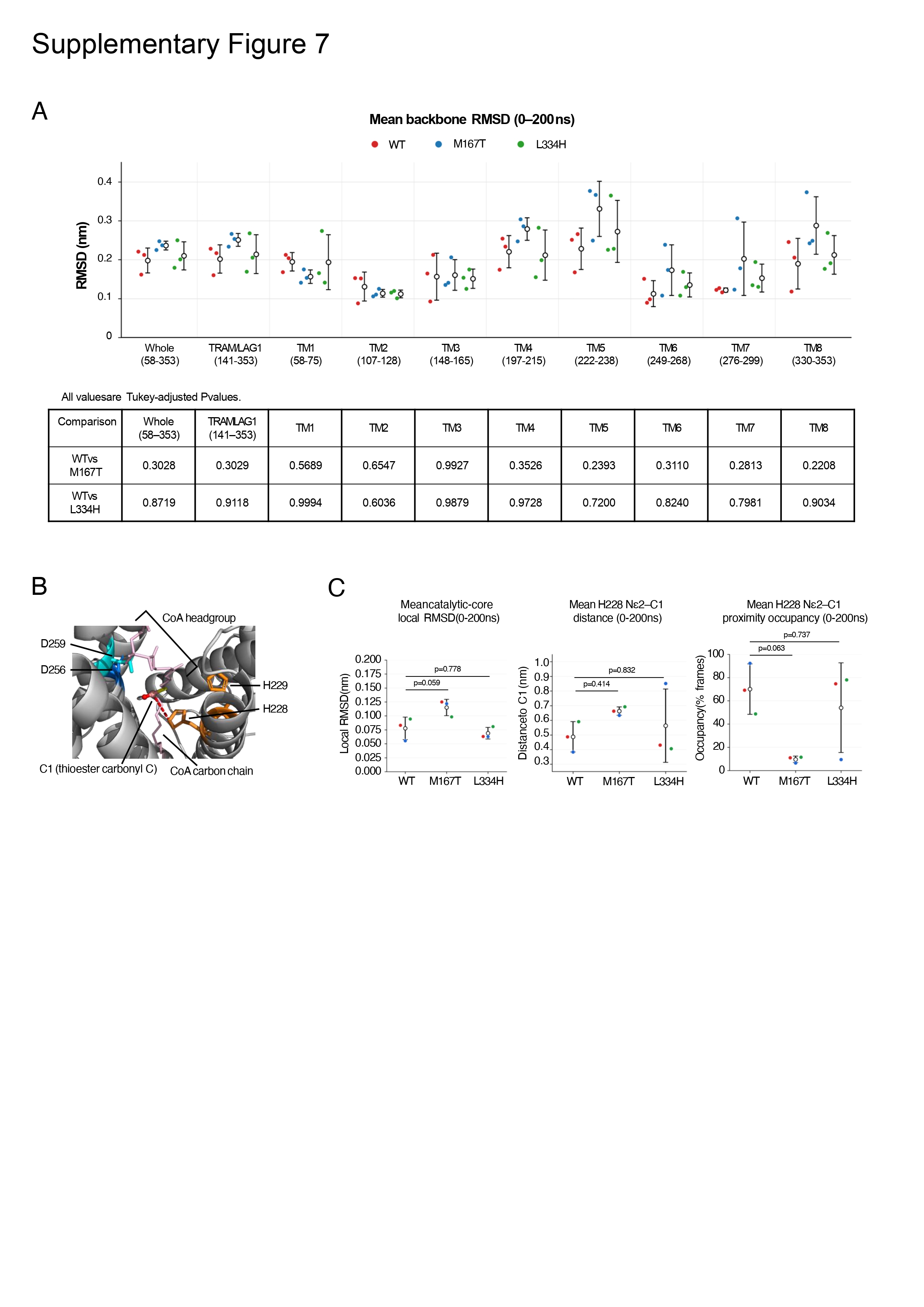
